# EIF3H Sustains Translational Programs Essential for Proliferation in Human Primed Pluripotency

**DOI:** 10.64898/2026.05.18.725809

**Authors:** Satoshi Suzuki, Chikako Okubo, Michiko Nakamura, Mari Hamao, Qi Fang, Knut Woltjen, Kazutoshi Takahashi

## Abstract

Pluripotent stem cells (PSCs) have remarkable capacity for unlimited self-renewal and differentiation into all somatic lineages. Although translational regulation has been implicated in the maintenance of PSC identity, the specific mechanisms involved remain poorly understood. Here, we identified EIF3H, a conserved subunit of the eIF3 translation initiation complex, as an essential regulator of human primed PSC proliferation and differentiation. CRISPR interference-mediated knockdown of EIF3H markedly reduced colony size, impaired proliferation, and diminished differentiation potential in all three germ layers. Integrated transcriptomic and translatomic profiling revealed that EIF3H loss decreased the translation of metal ion-related genes. Notably, the targeted suppression of metallothionein genes encoding metal-binding proteins recapitulated the proliferative defects observed in EIF3H-deficient PSCs, demonstrating a functional requirement for EIF3H-mediated translation of this gene family. Taken together, these findings establish EIF3H as a critical translational regulator that sustains PSC self-renewal and differentiation by maintaining the expression of key metabolic and stress-response genes, providing new insights into the molecular basis of pluripotency.

## INTRODUCTION

Pluripotent stem cells (PSCs) are defined by their unique capacity for unlimited self-renewal and potential to differentiate into all somatic lineages (1–5). These dual properties not only underpin their fundamental value in developmental biology but also establish PSCs as a cornerstone of regenerative medicine, enabling the generation of genetically modified models and renewable sources of cell-based therapeutics. Fully leveraging these capacities requires a deep mechanistic understanding of the molecular pathways that safeguard PSC identity and proliferative potential.

PSCs rely heavily on extrinsic cues for their maintenance, particularly the presence of growth factors in the culture environment. In primed human PSCs, the combination of basic fibroblast growth factor (bFGF) and transforming growth factor beta (TGF-β) is essential to sustain self-renewal and suppress spontaneous differentiation (6–8). These growth factors activate a network of intracellular signaling pathways that are further reinforced by interactions with adhesion molecules and extracellular matrix components. However, these signaling cascades are exquisitely balanced; even subtle perturbations can disrupt homeostasis and trigger differentiation, highlighting the necessity for tightly regulated extrinsic signaling.

In addition to extrinsic inputs, intrinsic cellular mechanisms are equally critical for maintaining PSC identity. Emerging evidence indicates that intracellular homeostasis is a key determinant of stem cell fate, particularly considering the frequent appearance of spontaneously differentiating cells under optimal culture conditions. These observations underscore the importance of the intrinsic regulatory networks that cooperate with extracellular cues to maintain self-renewal and pluripotency.

Among these intrinsic mechanisms, translational control has emerged as a central regulator of stem cell homeostasis (9). Translational regulation offers a rapid and energy-efficient mechanism for remodeling cellular states by selectively modulating protein synthesis from defined mRNA subsets (10). Consistent with this view, several translation-associated factors—including eIF4G2, RBPMS, and HTATSF1—have been shown to be indispensable for the differentiation capacity of human PSCs (11–14). Despite the increasing recognition of its importance, the roles of individual translation factors in establishing and maintaining PSC identity remain insufficiently understood.

To address this gap, we previously performed a genome-wide CRISPR interference (CRISPRi) screen and identified multiple components of the translation initiation machinery as essential regulators of PSC maintenance (15–17). In particular, we found that the loss of EIF3D, a subunit of the eukaryotic translation initiation factor 3 (eIF3) complex, led to the rapid collapse of pluripotency in primed PSCs. Given that the eIF3 complex coordinates one of the earliest steps of translation initiation and that several of its subunits have been implicated in selective mRNA recruitment, these findings raise the possibility that individual eIF3 components may perform subunit-specific regulatory functions in PSCs.

This notion was further supported by our screening, which revealed that EIF3H, a conserved core subunit of the eIF3 complex, is similarly required for PSC self-renewal (15). Intriguingly, previous studies on somatic cell types, such as HeLa cells, have reported only modest effects of EIF3H depletion on global translation and proliferation (18). These contrasting observations led us to hypothesize that EIF3H performs a function that is uniquely critical for the maintenance of human PSCs.

In the present study, we identify EIF3H as a previously unappreciated regulator of human PSC self-renewal and pluripotency. EIF3H depletion triggers severe proliferative retardation accompanied by weakened PSC identity, indicating its essential role in maintaining the PSC state. Remarkably, targeted suppression of metallothionein (MT) genes phenocopies the growth defects observed in EIF3H-deficient PSCs, establishing a functional link between EIF3H-dependent translation and PSC fitness. Together, these findings reveal a hitherto unrecognized translational module that is indispensable for sustaining both the identity and proliferative capacity of human PSCs.

## RESULTS

### EIF3H Is essential for the self-renewal of human PSCs

To elucidate the role of EIF3H in human PSCs, we established doxycycline (Dox)-inducible CRISPRi knockdown (KD) lines targeting EIF3H. For comparison, we used three independent cell lines expressing non-targeting single-guide RNA (sgRNA) (15–17). Unless otherwise indicated, all experiments were conducted using three independently derived clonal lines per treatment. Quantitative reverse transcription-polymerase chain reaction (RT-PCR) analyses confirmed that Dox treatment for three days led to >99% suppression of *EIF3H* transcripts, and this high KD efficiency was sustained for at least 12 days (Fig. 1A).

**Fig. 1:**
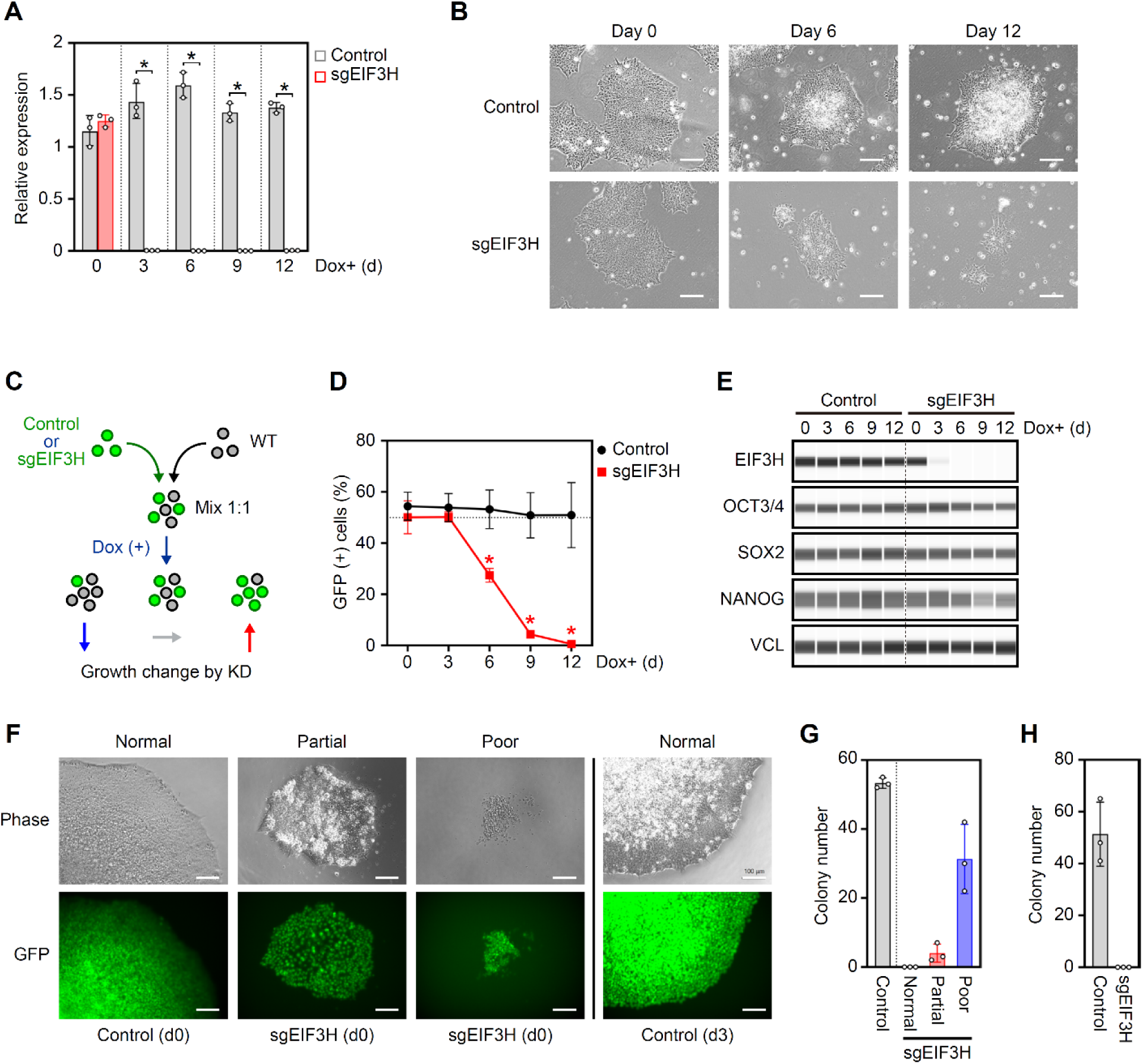
EIF3H is essential for PSC self-renewal. **(A),** RNA expression of EIF3H on the indicated days after Dox addition (mean ± SD, n=3). Values were normalized to *GAPDH* and compared with the control without Dox. P-values were determined using an unpaired t-test: P_d3_=0.0048; P_d6_=0.0020; P_d9_=0.0014; P_d12_=0.0004. **(B),** Representative images of control and EIF3H KD PSCs on days 0, 6, and 12 post-KD induction. The cells were passaged every three days; therefore, the images show the colonies three days after plating. Scale bars: 100 μm. **(C),** The scheme of growth competition assay. **(D),** Percentages of GFP-positive cells on indicated days in the competition assay (mean ± SD, n=3). P-values were determined using an unpaired t-test: P_d6_=0.0044; P_d9_=0.0034; P_d12_=0.0007 **(E),** Expression of pluripotency-associated proteins during EIF3H KD. VINCULIN (VCL) was used as the loading control. **(F),** Representative images of single-cell-derived control and EIF3H KD colonies plated before and 3 days after Dox addition. Scale bars: 100 μm. **(G),** Number of wells containing normal, partial, and poor grown colonies derived from cells plated before Dox addition (mean ± SD, n=3). **(H),** Number of wells containing colonies derived from cells plated on day 3 after Dox addition (mean ± SD, n=3).

Six days after KD, phenotypic observations revealed no obvious changes in cell morphology, but a tendency toward a smaller size compared to the control was observed (Fig. 1B). By day 12, the EIF3H KD colonies had become markedly smaller and more compact. They displayed sharp and irregular boundaries, suggesting disruptions in intercellular adhesion or proliferation kinetics (Fig. 1B). In contrast, the control PSCs retained the hallmark features of undifferentiated stem cell colonies. These observations implicate EIF3H is a key factor in maintaining the colony architecture and proliferative potential of human PSCs.

To quantitatively verify the results of cell observation, we performed a competitive proliferation assay. Wild-type (green fluorescent protein [GFP]-negative) PSCs were co-cultured in a 1:1 ratio with either GFP-positive control or GFP-positive EIF3H KD cells (Fig. 1C). In the presence of Dox, the fraction of GFP-positive EIF3H KD cells began to considerably decline from day 6 onward, whereas the ratio remained stable in control co-cultures (Fig. 1D). This loss was not compensated by co-cultured wild-type cells, indicating that the EIF3H depletion phenotype is cell-autonomous.

To explore the malfunction of the highly proliferative properties of PSCs, we examined the expression of core PSC markers in EIF3H KD cells over time (Fig. 1E). The results showed that OCT3/4, SOX2, and NANOG expression did not substantially decrease until 6 days after KD induction. However, a decrease in NANOG expression was observed on day 9 after the addition of Dox. These results suggest that the cell proliferation defects caused by EIF3H KD occur prior to changes in PSC characteristics, including colony morphology and PSC marker expression.

To examine cell dynamics in more detail, we conducted a clonal colony formation assay by plating PSCs at single-cell density onto a 96-well plate either before (Day 0) or after (Day 3) Dox treatment and assessing colony growth under continuous Dox exposure. In the case of plating before Dox addition, a few wells showed proliferation of EIF3H KD colonies. However, they were smaller than those of the control PSCs, and GFP fluorescence images revealed an uneven cell density in these colonies. Colonies derived from EIF3H KD PSCs observed in a relatively large number of wells were abnormal and contained few cells. The control PSCs did not form such abnormal colonies. EIF3H KD cells consistently generated fewer and smaller colonies than control cells (Figs. 1F and 1G). After Dox addition, plated EIF3H KD PSCs were unable to form colonies, which coincided with maximal EIF3H suppression (Fig. 1H).

Collectively, these results establish EIF3H as an essential regulator of human PSC self-renewal, acting through the maintenance of proliferative capacity and colony-forming potential under defined culture conditions.

### EIF3H sustains the proliferative capacity that underlies stable pluripotency

The results presented above suggest that EIF3H depletion leads to abnormalities in the expansion of clonal and bulk PSC populations. However, the underlying cause remains unclear. Therefore, we next sought to determine whether the observed reduction in EIF3H KD cell numbers could be attributed to impaired adhesion to the culture substrate. To this end, equal numbers of control and EIF3H KD cells were plated on days 3 or 5 following Dox addition, and the number of adherent living and dead cells was quantified 24 h later by observing GFP and 4′,6-diamidino-2-phenylindole (DAPI) fluorescence, respectively. There was no significant difference in the initial adhesion efficiency between control and EIF3H KD cells, indicating that defects in cell adhesion are unlikely to account for the decreased cell numbers in EIF3H-depleted PSCs (Fig. 2A).

**Fig. 2:**
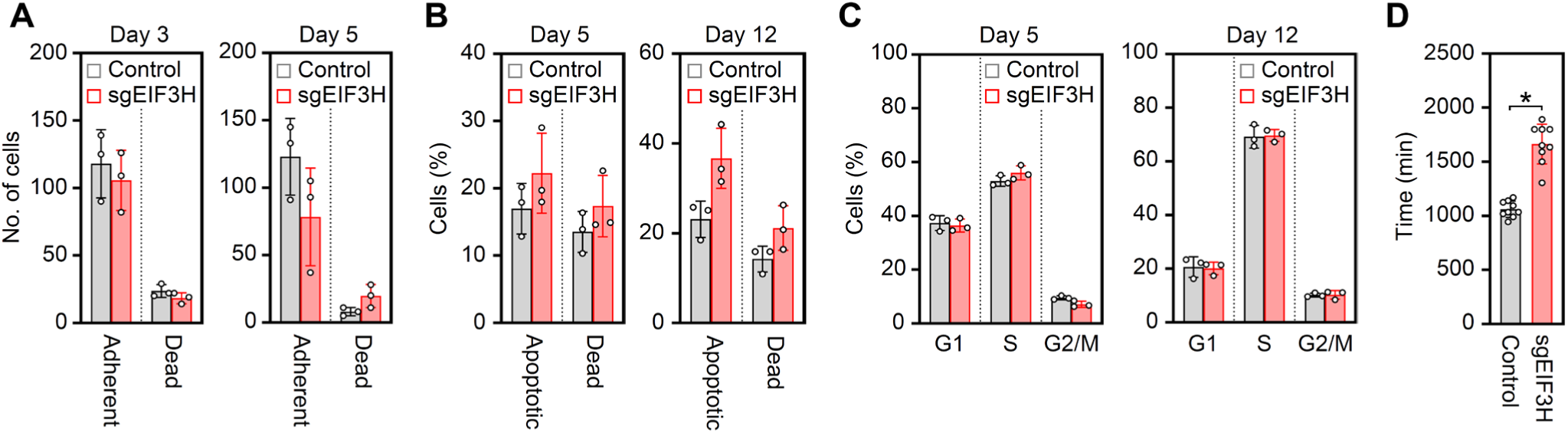
Deficiency of EIF3H causes proliferation delay in PSCs. **(A),** Adhesion efficiency of control and EIF3H KD PSCs plated on days 3 (left) and 5 (right) after Dox addition (mean ± SD, n=3). P-values were determined using an unpaired t-test. **(B),** Percentages of apoptotic and dead cells in control and EIF3H KD PSCs (mean ± SD, n=3). P-values were determined using an unpaired t-test. **(C),** Cell cycle phase distribution of control and EIF3H KD PSCs on days 5 (left) and 12 (right) after doxycycline addition (mean ± SD, n=3). No significant differences were observed in any cell cycle phase on either day 5 or day 12. P-values were calculated using an unpaired t-test. **(D),** Time required for one cell cycle of control and EIF3H KD PSCs on day 5 after Dox addition (mean ± SD, n=9). P-values were determined using an unpaired t-test: P=2.4294E-06.

To examine whether EIF3H depletion induced cell death, we performed Annexin V staining to assess the apoptotic cell populations. Across multiple biological replicates, no significant increase in apoptosis was detected in EIF3H KD cells compared to controls on days 5 or 12 post-KD induction (Fig. 2B). This suggests that EIF3H is not primarily involved in cell survival pathways and that apoptosis and aberrant cell death do not account for the proliferative defect observed.

Subsequently, we evaluated whether the reduction in cell numbers resulted from cell cycle alterations. 5-ethynyl-2’-deoxyuridine (EdU) incorporation assays and cell cycle content analysis were performed on day 5 post-Dox addition, corresponding to the early phase of observed cell number decline. Surprisingly, no substantial changes were detected in cell cycle phase distribution at this early time point (Fig. 2C). However, by day 12 post-induction, EIF3H KD cells exhibited a marked increase in the G1 population, indicating delayed progression through the G1/S checkpoint (Fig. 2C). This shift, while notable, occurred after the initial reduction in cell numbers and therefore does not explain the early proliferative deficit.

To gain further insight into the cell cycle dynamics of EIF3H KD cells, we utilized human PSCs stably expressing the tandem fluorescent ubiquitination-based cell cycle indicator (tFucci) system, a fluorescent cell cycle reporter that distinguishes the G1, S, and G2/M phases in living cells (19). Cells were seeded on day 5 post Dox addition and subjected to 48 h of time-lapse imaging. Control PSCs completed a full cell cycle within approximately 1,000 min, which is consistent with previously published values (20) (Fig. 2D). In contrast, EIF3H KD cells required approximately 1,700 min to complete single-cell division. The observation that EIF3H KD cells require 1.7 times longer to complete a single cell cycle compared to control cells aligns with the reduced proliferation rate of EIF3H KD cells in the growth competition experiment (Fig. 1D). These results provide compelling evidence that EIF3H depletion leads to a marked prolongation of the cell cycle rather than arrest at a specific phase. Importantly, this cell cycle lengthening precedes visible morphological changes or the reduction of PSC markers, suggesting that EIF3H plays a direct role in sustaining the rapid proliferative kinetic characteristics of primed human PSCs (Figs. 1B and 1C). Taken together, our results demonstrate that EIF3H is indispensable for the maintenance of self-renewal in primed human PSCs. EIF3H depletion leads to reduced colony size, impaired clonal expansion, and a progressive decline in cell number, all in the absence of increased apoptosis or adhesion defects. The primary mechanism underlying these phenotypes appears to be a significant extension of the cell cycle, which ultimately compromises the proliferative capacity of PSCs.

### Loss of EIF3H perturbs gene expression in PSCs

To gain a comprehensive understanding of the molecular consequences of EIF3H depletion in human PSCs, we performed RNA sequencing (RNA-seq) at multiple time points (days 3, 5, and 12) following Dox addition. Comparative analysis of gene expression between the control and EIF3H KD revealed a progressive increase in the number of differentially expressed genes (DEGs) over time post-KD induction (Figs. 3A and 3B). By day 5, prominent downregulation of the *MT* gene cluster was observed, a trend that became even more pronounced by day 12 (Figs. 3C and S1). Gene ontology (GO) analysis revealed dynamic and time-dependent changes in gene expression following EIF3H KD. On day 3, the upregulated genes in EIF3H KD cells were enriched for categories related to small nuclear ribonucleoproteins (snRNPs) and differentiation, whereas the downregulated genes were not enriched for any specific functional category (Table S1). By day 5, the upregulated genes were predominantly associated with neural differentiation and cell adhesion, as the changes on day 3 were more pronounced, whereas the downregulated genes showed enrichment for term related to cell growth (Tables S2 and S3). Notably, the expression patterns observed on day 12 closely mirrored those on day 5, with similar enrichment profiles in both upregulated and downregulated gene sets, suggesting that the transcriptional impact of EIF3H depletion stabilizes by this time point (Tables S4 and S5).

**Fig. 3:**
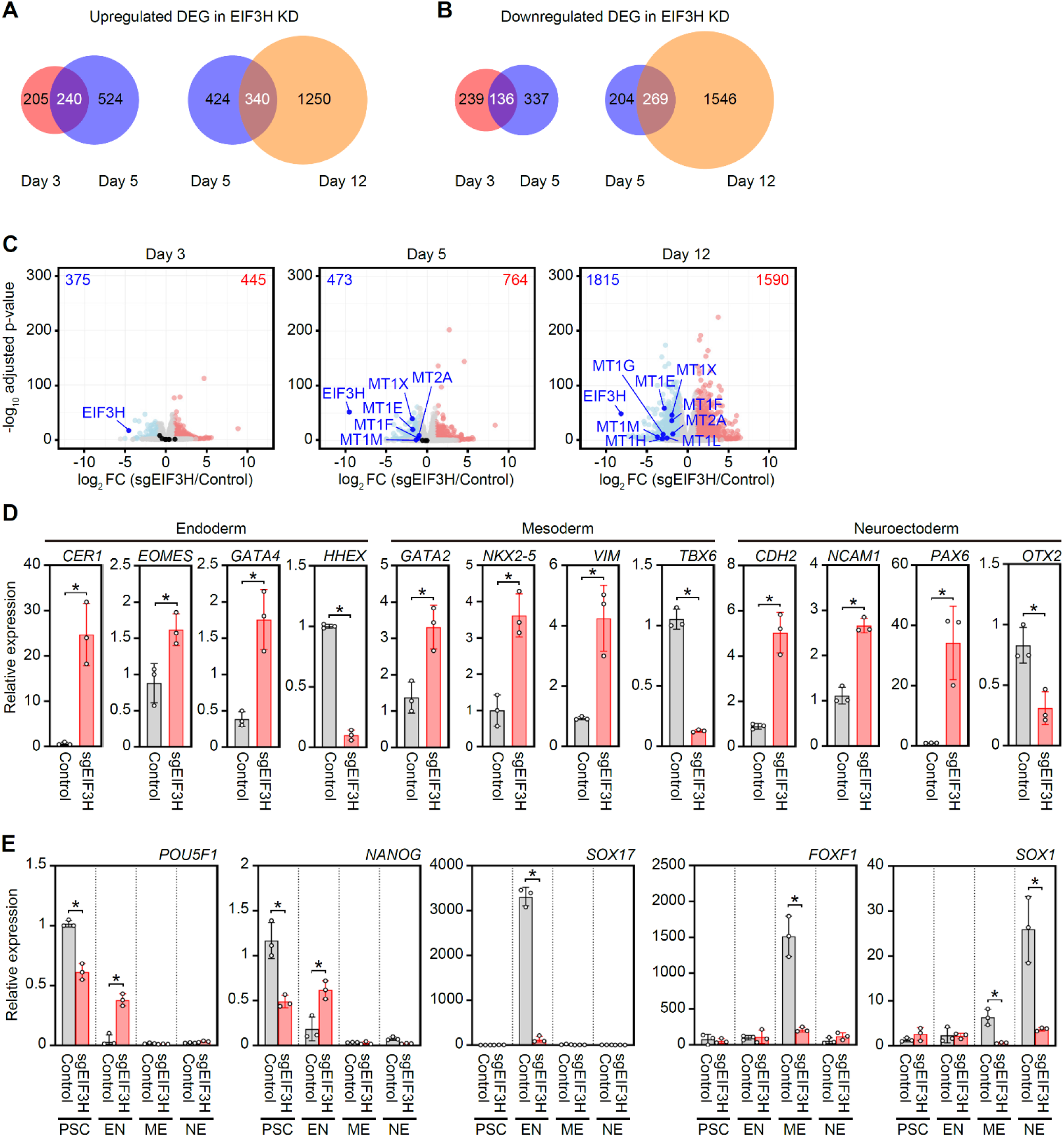
Loss of EIF3H compromises differentiation capacity of human PSCs. **(A),** Venn diagram illustrating the overlap of upregulated DEGs between days 3 and 5 (left) and days 5 and 12 (right) following EIF3H KD. **(B),** Venn diagram illustrating the overlap of downregulated DEGs between days 3 and 5 (left) and days 5 and 12 (right) following EIF3H KD. **(C),** Volcano plots showing DEGs between control and sgEIF3H PSCs on days 3, 5, and 12 after Dox addition. The highlighted genes indicate EIF3H and MT family genes. **(D),** Relative expression of selected lineage-associated marker genes on day 12 post-KD induction (mean ± SD, n=3). P-values were calculated using an unpaired t-test: P_CER1_=0.0258; P_EOMES_=0.0229; P_GATA4_=0.0004; P_HHEX_=0.0002; P_GATA2_=0.0130; P_NKX2-5_=0.0047; P_VIM_=0.0341; P_TBX6_=0.0023; P_CDH2_=0.0142; P_NCAM1_=0.0004; P_PAX6_=0.0421; P_OTX2_=0.0112. **(E),** RNA expression of pluripotency and lineage marker genes in undifferentiated PSCs and directed differentiation derivatives, including endoderm (EN), mesoderm (ME), and neuroectoderm (NE), normalized to *GAPDH* and compared with control PSCs (mean ± SD, n=3). P-values were calculated using an unpaired t-test: *POU5F1* (P_PSC_=0.00385; P_EN_=0.0012); *NANOG* (P_PSC_=0.0191; P_EN_=0.0122; P_ME_=0.0343); *SOX17* (P_EN_=0.000423); *FOXF1* (P_ME_=0.0143); *SOX1* (P_ME_=0.0299; P_NE_=0.0339). No significant differences were observed in the other comparisons.

The impact of EIF3H KD on the core pluripotency network was relatively modest. Among the 19 canonical PSC marker genes examined (21–25), only six (*NANOG*, *DPPA2*, *IFITM1*, *FGF4*, *GDF3*, and *UTF1*) were significantly downregulated by day 12 compared with the controls under the criteria of fold change (FC) > 2 and p < 0.05 (Table S6). These results suggest that EIF3H KD has a limited impact on the pluripotency maintenance network.

In contrast, EIF3H KD substantially dysregulated the genes associated with early lineage commitment (Fig. 3D). These results are consistent with the enrichment of terms related to cell differentiation, including the nervous system, among the genes whose expression increased with EIF3H KD. This suggests the acquisition of aberrant differentiation-like transcriptional signatures, despite the incomplete loss of the undifferentiated state. Indeed, directed differentiation toward each germ layer revealed diminished activation of lineage markers such as *SOX17* (endoderm: EN), *FOXF1* (mesoderm: ME), and *SOX1* (neuroectoderm: NE) in EIF3H KD cells (Fig. 3E). These results demonstrate that EIF3H is necessary for the precise differentiation of human PSCs into all three germ layers.

Together, these findings demonstrate that EIF3H depletion perturbs the gene expression landscape of human PSCs, which is characterized by the suppression of MT-related genes and perturbation of pluripotency-associated transcripts and differentiation markers.

### EIF3H regulates the translation of specific mRNAs

To determine whether EIF3H directly contributed to translational control in PSCs, we assessed global protein synthesis. EIF3H KD modestly, but reproducibly, reduced puromycin incorporation (84.4% of the control; Fig. 4A) and induced a pronounced accumulation of 80S monosomes (Fig. 4B). These findings indicate that EIF3H loss perturbs translation initiation—a defect consistent with EIF3H functioning within the eIF3 complex (15, 26).

**Fig. 4:**
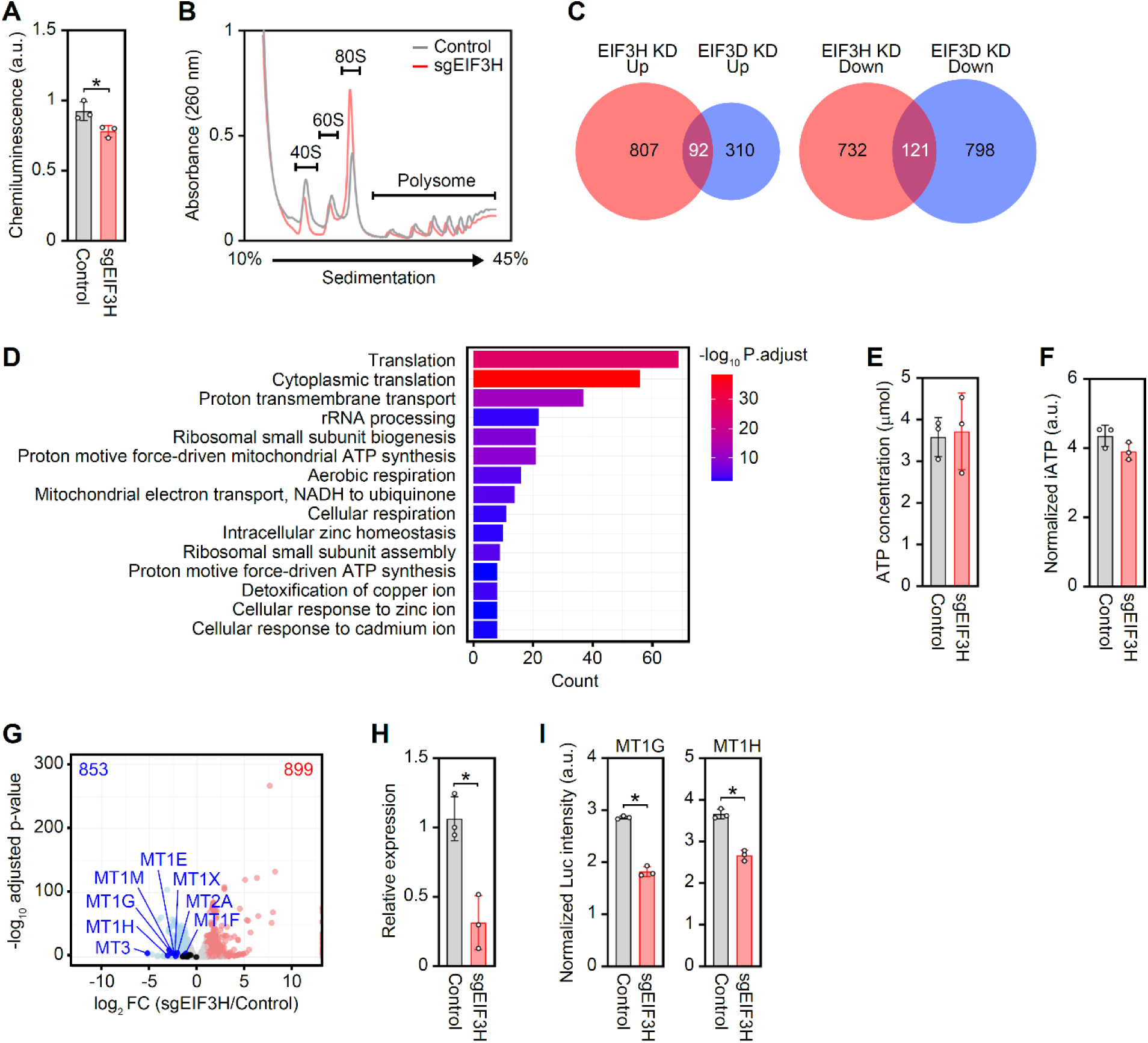
Translational dysregulation of *MT* genes following EIF3H KD. **(A),** Quantification of *de novo* protein synthesis by measuring puromycin incorporation in control and sgEIF3H PSCs on day 3 after Dox induction. Puromycin signals were normalized to those of control cells (mean ± SD, n=3). P-values were calculated using an unpaired t-test: P=0.0427. **(B),** Representative polysome profiles of control (black) and sgEIF3H (red) PSCs on day 3 after doxycycline (Dox) induction. Ribosomal fractions corresponding to 40S, 60S, 80S, and polysomes are indicated. **(C),** Venn diagrams showing the overlap of genes with increased (left) or decreased (right) translation efficiency (TE) in EIF3H KD and EIF3D KD PSCs. **(D),** Gene ontology (GO) enrichment analysis of genes with decreased translation efficiency (TE) following EIF3H KD. Bar plot showing significantly enriched GO terms (top 15) among TE-downregulated genes identified by ribosome profiling on day 3 after doxycycline (Dox) induction. The x-axis indicates the number of genes associated with each GO term (Count), and the color scale represents the significance level (−log_10_ adjusted P-value). Enriched categories include translation-, mitochondrial-, and metal ion homeostasis–related processes. **(E),** Quantification of intracellular ATP levels in control and sgEIF3H PSCs using a luminescence-based ATP assay. ATP concentrations were calculated from a standard curve and are shown as mean ± SD, n=3. P-values were determined using an unpaired t-test. **(F),** Quantification of normalized mitochondrial ATP signals in control and sgEIF3H PSCs on day 5 after doxycycline (Dox) induction. ATP signals were measured by flow cytometry using synthesized mRNA encoded ATP sensor mito-iATPSnFR2. Mean ± SD, n = 3 independent clones. P-values were determined using an unpaired t-test. **(G),** Volcano plot showing differentially translated genes in EIF3H KD PSCs on day 3 after Dox induction based on Ribo-seq analysis. The x-axis indicates log_2_ fold change of translation efficiency (sgEIF3H/control), and the y-axis indicates −log_10_ adjusted P-value. Metallothionein (MT) family genes are highlighted in blue. **(H),** Quantification of MT protein levels by ELISA in control and sgEIF3H PSCs (mean ± SD, n=3). P-values were calculated using an unpaired t-test: P=0.0075. **(I),** Luciferase reporter assays for *MT1G* and *MT1H* 5′UTRs in control and sgEIF3H PSCs. Normalized luciferase intensities are shown (mean ± SD, n=3). P-values were calculated using an unpaired t-test: P_MT1G_=0.0014; P_MT1H_=0.0006.

Ribosome profiling on day 3, prior to overt phenotypic changes (Figs. 1A, 1D, and 1E) revealed widespread translational rewiring. Translation efficiency (TE) increased for 899 genes and decreased for 853 genes (Fig. S2). Notably, the large number of TE-upregulated genes contrasts sharply with our previous results for EIF3D depletion (15), indicating subunit-specific translational programs within the eIF3 complex (Fig. 4C). GO analysis of TE-upregulated genes enriched transcription-, chromatin-, and RNA processing–related categories (Table S7), further underscoring the unique regulatory function of EIF3H in PSCs.

EIF3H KD did not substantially alter the TE of core pluripotency genes, suggesting that the attenuation of pluripotency-associated gene expression may be a secondary effect caused by reduced proliferation. Among the TE-downregulated genes, GO terms related to translation, mitochondria, and metal-ion metabolism were enriched (Fig. 4 and Table S8). Although numerous mitochondria-associated transcripts showed reduced TE, intracellular ATP levels measured by a luminescence-based assay (Fig. 4E) and mitochondrial ATP signals (Fig. 4F) measured using mito-iATPSnFR2 (27) remained unchanged, suggesting that overt ATP depletion is unlikely to explain the EIF3H KD phenotype.

*MT* genes were among the most strongly affected targets. Further, MT translation efficiency decreased markedly as early as day 3 (Fig. 4G), whereas RNA levels did not decline until day 5 (Fig. 3C). This clear temporal separation suggests that impaired translation is the initiating event and that secondary transcript downregulation occurs. Enzyme-linked immunosorbent assay (ELISA) revealed that MT protein levels were markedly reduced by EIF3H KD (Fig. 4H). Therefore, to test whether *MT* family genes are direct translational targets of EIF3H, we used luciferase reporters containing *MT1G* or *MT1H* 5′UTRs. Both reporters exhibited reduced activity upon EIF3H KD, indicating 5′UTR-dependent translational regulation (Fig. 4I). Together with the early decrease in TE, these findings establish MT as the key EIF3H-dependent translational targets required for PSC homeostasis.

These findings position MT as the earliest and most sensitive translational targets of EIF3H, defining impaired MT translation as the primary molecular defect preceding downstream transcriptomic alterations.

### MT, a target of EIF3H, promotes PSC proliferation

Transcriptomic and translatomic analyses suggested that MTs are major EIF3H targets. To test their functional contributions to PSC self-renewal, we sought to suppress MT expression. However, because humans possess 11 *MT* genes with substantial redundancy, we knocked down metal regulatory transcription factor 1 (MTF1), a transcription factor that regulates *MT* family expression (28, 29) (Fig. 5A). RNA-seq confirmed broad *MT* downregulation in MTF1 KD PSCs (Fig. 5B). GO analysis of MTF1 KD DEGs showed that downregulated genes were enriched for metal ion transport and metal-response pathways (Table S9). Genes downregulated in both EIF3H KD and MTF1 KD significantly overlapped, particularly within the metal-ion response pathways (Fig. 5C and Table S10). MT protein expression did not significantly decrease compared to that in the control on day 3 of MTF1 KD; however, it decreased markedly by day 5. This trend was similar to that observed for EIF3H KD (Fig. 5D). Functionally, MTF1 KD cells progressively lost cell growth beginning around day 6 and showed a considerable decrease in cell population by day 9. The timing of decreased cell growth in MTF1 KD was slightly slower than that in EIF3H KD but mirrored the proliferative decline observed in EIF3H KD cells (Fig. 5E). Similar to EIF3H loss, MTF1 KD did not alter cell cycle distribution or induce apoptosis (Figs. 5G and 5H), suggesting a shared cell-autonomous mechanism.

**Fig. 5:**
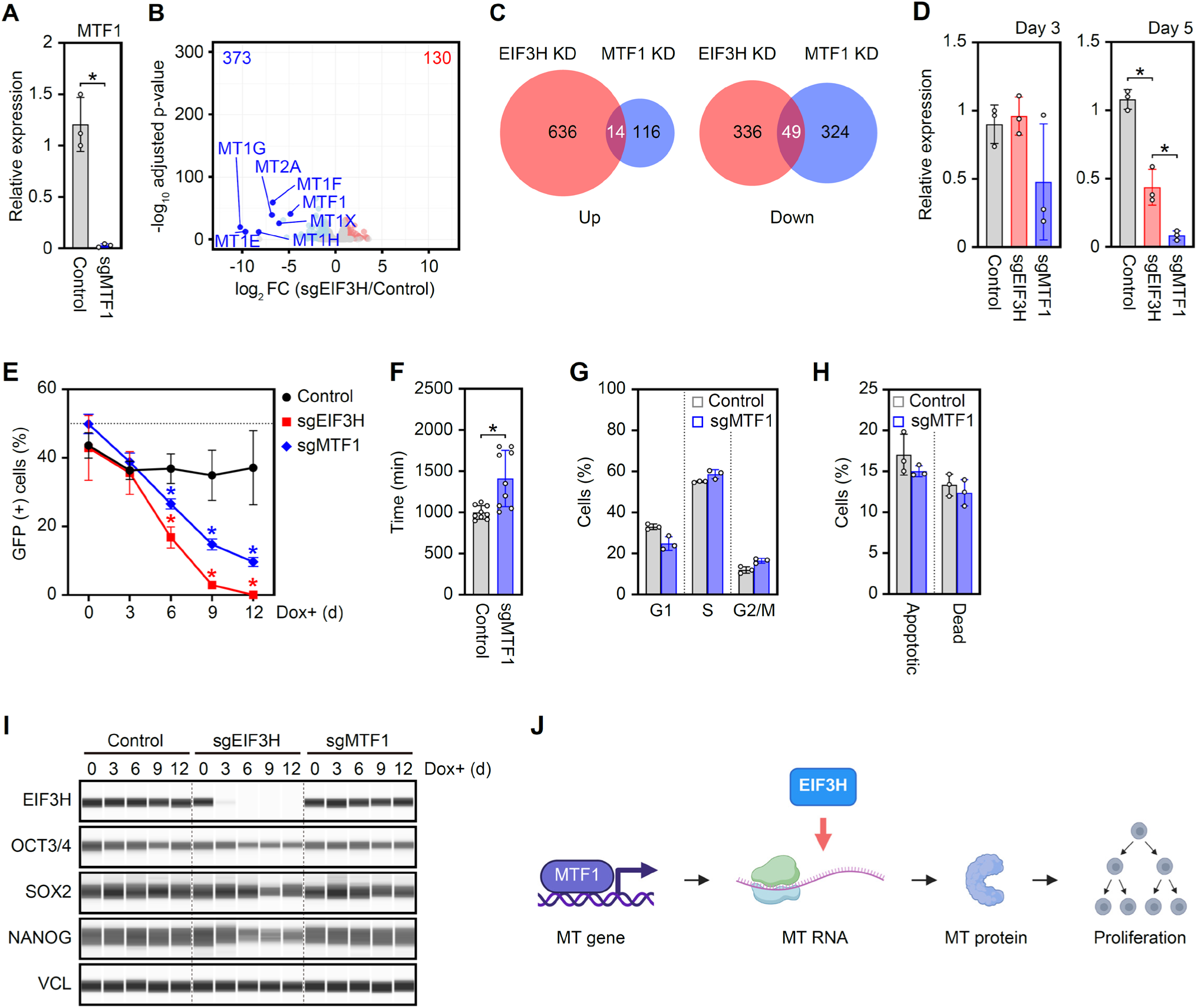
MTs act downstream of EIF3H to support proliferation of human PSCs. **(A),** Quantitative PCR analysis of *MTF1* mRNA expression in control and sgMTF1 PSCs on day 5 after doxycycline (Dox) induction. Expression levels were normalized to *GAPDH* and expressed relative to control cells (mean ± SD, n=3). P-values were calculated using an unpaired t-test: P=0.0158. **(B),** Volcano plot showing differentially expressed genes between control and sgMTF1 PSCs on day 5 after Dox induction based on RNA-seq analysis. The x-axis indicates log₂ fold change (sgMTF1/control), and the y-axis indicates−log_10_ adjusted p-value. MT family genes are highlighted in blue. **(C),** Venn diagrams showing the overlap of differentially expressed genes identified by RNA-seq between sgEIF3H and sgMTF1 PSCs on day 5 after Dox induction. **(D),** Quantification of MT protein levels measured by ELISA in control, sgEIF3H, and sgMTF1 PSCs on days 3 and 5 after Dox induction (mean ± SD, n=3). P-values were calculated using an unpaired t-test. On day 3, no significant differences were observed between the groups. On day 5, P-values were as follows: control vs. sgEIF3H (P=0.0045); control vs. sgMTF1 (=0.0003); sgEIF3H vs. sgMTF1 (P=0.0360). **(E),** Growth competition assay of control, sgEIF3H, and sgMTF1 PSCs. GFP-positive cell fractions were quantified by flow cytometry at the indicated time points following Dox induction (mean ± SD, n=3). P-values were calculated using an unpaired t-test: control vs. sgEIF3H (P_d6_=0.0028; P_d9_=0.0016; P_d12_=0.0040); control vs. sgMTF1 (P_d6_=0.0175; P_d9_=0.00948; P_d12_=0.0119). **(F),** Quantification of cell cycle duration measured by live-cell imaging using tFucci-expressing sgMTF1 PSCs starting on day 7 after Dox induction (mean ± SD, n=3). P-values were calculated using an unpaired t-test: P=0.0067. **(G),** Cell cycle phase distribution of control and sgMTF1 PSCs on day 5 after Dox induction as determined by EdU incorporation and DNA content analysis (mean ± SD, n=3). P-values were determined using an unpaired t-test. **(H),** Quantification of apoptotic and dead cell populations in control and sgMTF1 PSCs on day 5 after Dox induction was performed using Annexin V and SYTOX Blue staining (mean ± SD, n=3). P-values were determined using an unpaired t-test. **(I),** Expression of EIF3H and pluripotency-associated proteins in control, sgEIF3H, and sgMTF1 PSCs on the indicated days after Dox induction. VINCULIN (VCL) was used as the loading control. **(J),** Schematic model illustrating the proposed EIF3H–MT axis underlying the regulation of human PSC proliferation and self-renewal.

EIF3H KD gradually reduced PSC marker expression, whereas MTF1 KD did not affect the PSC marker expression (Fig. 5I). These results suggest that MTF1 KD-induced MT depletion contributes specifically to proliferative capacity rather than to the maintenance of the undifferentiated state. Based on these results, we concluded that EIF3H-mediated translation promotion of *MT* transcripts enhances the proliferative capacity of human PSCs by increasing MT protein expression.

These findings demonstrate that MTs constitute major effectors downstream of EIF3H. Combined with the early TE reduction observed in Fig. 4, impaired MT translation was established as the primary initiating defect in EIF3H-deficient PSCs. Therefore, this study establishes a model of human PSC self-renewal driven by the EIF3H-MT axis-mediated proliferation (Fig. 5J).

## DISCUSSION

In this study, we identified EIF3H as a critical regulator of self-renewal in human PSCs. Loss of EIF3H led to a marked decline in cellular proliferation beginning on day 3 post-KD, which was followed by a gradual reduction in the expression of key pluripotency markers from day 6. Importantly, at the early stage of the proliferation decline (day 3), we observed a coordinated decrease in the translation of genes involved in core cellular functions, including those related to translation, mitochondrial activity, and metal ion homeostasis. These findings suggest that EIF3H supports PSC self-renewal by maintaining the translation of factors required for cellular homeostasis. Taken together, our results reveal a mechanistic link between EIF3H-mediated translational control and the maintenance of PSC identity and growth, highlighting EIF3H as a pivotal node in the preservation of pluripotent cell function.

In this study, we identified MT as a key translational target of EIF3H in human PSCs. While previous studies have reported that MT is highly expressed in undifferentiated PSCs and downregulated upon differentiation (30), the functional relevance of this observation has remained unclear. Previous studies in cancer cells have also implicated MTs in cell-cycle regulation and proliferation (31, 32), but whether MTs functionally support proliferation in human PSCs had remained unclear. To address this, we reduced MT expression by knocking down MTF1, a transcription factor that regulates *MT* gene expression. This perturbation led to a marked reduction in PSC proliferation, demonstrating that MT plays an active role in supporting the proliferative capacity of PSCs. Based on these findings, we propose a model in which EIF3H-mediated translational regulation ensures high levels of MT expression in undifferentiated PSCs, thereby contributing to the maintenance of their rapid proliferative state. This highlights a previously unknown link between translational control and metabolic or protective functions of MT in sustaining PSC self-renewal.

Recent studies have reported that EIF3H KD in HeLa cells does not significantly affect cell proliferation (18). Furthermore, the translational consequences of EIF3H depletion appear to be minimal in this context, with only 21 transcripts exhibiting altered translation efficiency. These findings contrast with our observations in PSCs, in which EIF3H plays a critical role in sustaining proliferation and regulating the translation of a broader set of genes. In contrast, it has been reported that EIF3H promotes the proliferation of some cancer cells, which aligns with our findings in PSCs (33–35). While differences in the experimental setup may account for some of these discrepancies, the results strongly suggest that the function of EIF3H is highly dependent on the cellular context. To further explore the specificity of the EIF3H function, we compared our EIF3H data with our previously published EIF3D data (15). Despite their physical association within the eIF3 complex, the two subunits play different functional roles. This divergence has also been noted in the aforementioned studies using HeLa cells (18), reinforcing the idea that EIF3H and EIF3D contribute to translational control in a manner specific to the subunit and cell type. In contrast, impaired differentiation potential was commonly observed in both EIF3H and EIF3D KD human PSCs, indicating that normal eIF3 complex functions are essential for differentiation (15). Overall, our findings highlight the unique and vital role of EIF3H in maintaining the translational landscape and proliferative capacity of PSCs.

In summary, this study demonstrates that translational control of specific gene sets by EIF3H plays a pivotal role in sustaining the high proliferative capacity of human PSCs. Our findings provide a coherent model that connects previously observed, yet unexplained, phenomena with the fundamental characteristics of PSCs. By shedding light on the underexplored layer of translational regulation, our work not only deepens our understanding of PSC biology but also opens new avenues for uncovering the molecular mechanisms that govern stem cell identity and function.

## ACKNOWLEDGEMENTS

We thank B. Conklin, M. Ikeya, M. Iwasaki, A. Miyawaki, Y. Sato, S. Yamanaka, F. Zhang, and CiRA Common Equipment Management Office for sharing materials and equipment; S. Kume, T. Murakami, T. Nakamura, L. Ni, A. Ogawa, N. Shiraki, and K. Tomoda for technical assistance and discussion; Y. Ishida and S. Takeshima for administrative support; and Editage for English language editing. The schematic figures were created in BioRender.com.

## AUTHOR CONTRIBUTIONS

Conceptualization: SS, CO, KT; Methodology: SS, CO, QF, KW, KT; Investigation: SS, CO, MN, MH, KT; Visualization: SS, CO, KT; Funding acquisition: SS, CO, KT; Project administration: CO, KT; Supervision: CO, KT; Writing – original draft: SS, CO, KT; Writing – review & editing: all authors

---

**Fig. S1:**
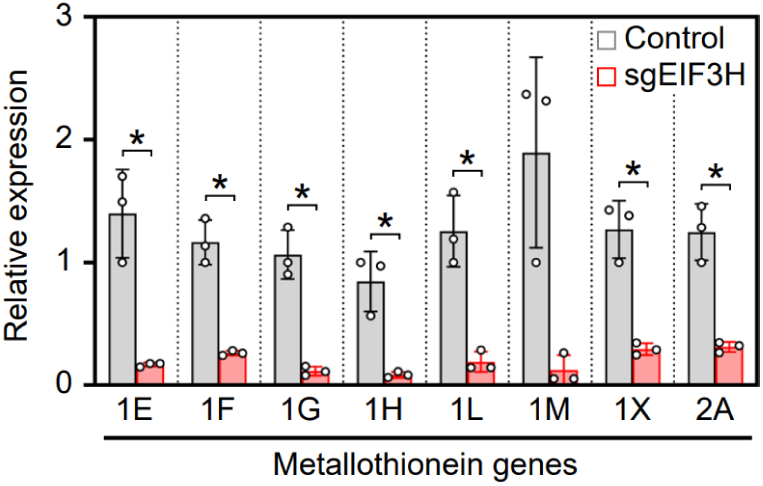
MT gene expression in EIF3H KD PSC (Related to Fig. 3) Relative expression of *MT* family genes in control and sgEIF3H PSCs on day 12 post-Dox addition derived from RNA-seq data (mean ± SD, n=3). P-values were determined by unpaired t-test: P_1E_=0.0271; P_1F_=0.0124; P_1G_=0.0122; P_1H_=0.0313; P_1L_=0.0179; P_1X_=0.0155; P_2A_=0.0172.

**Fig. S2:**
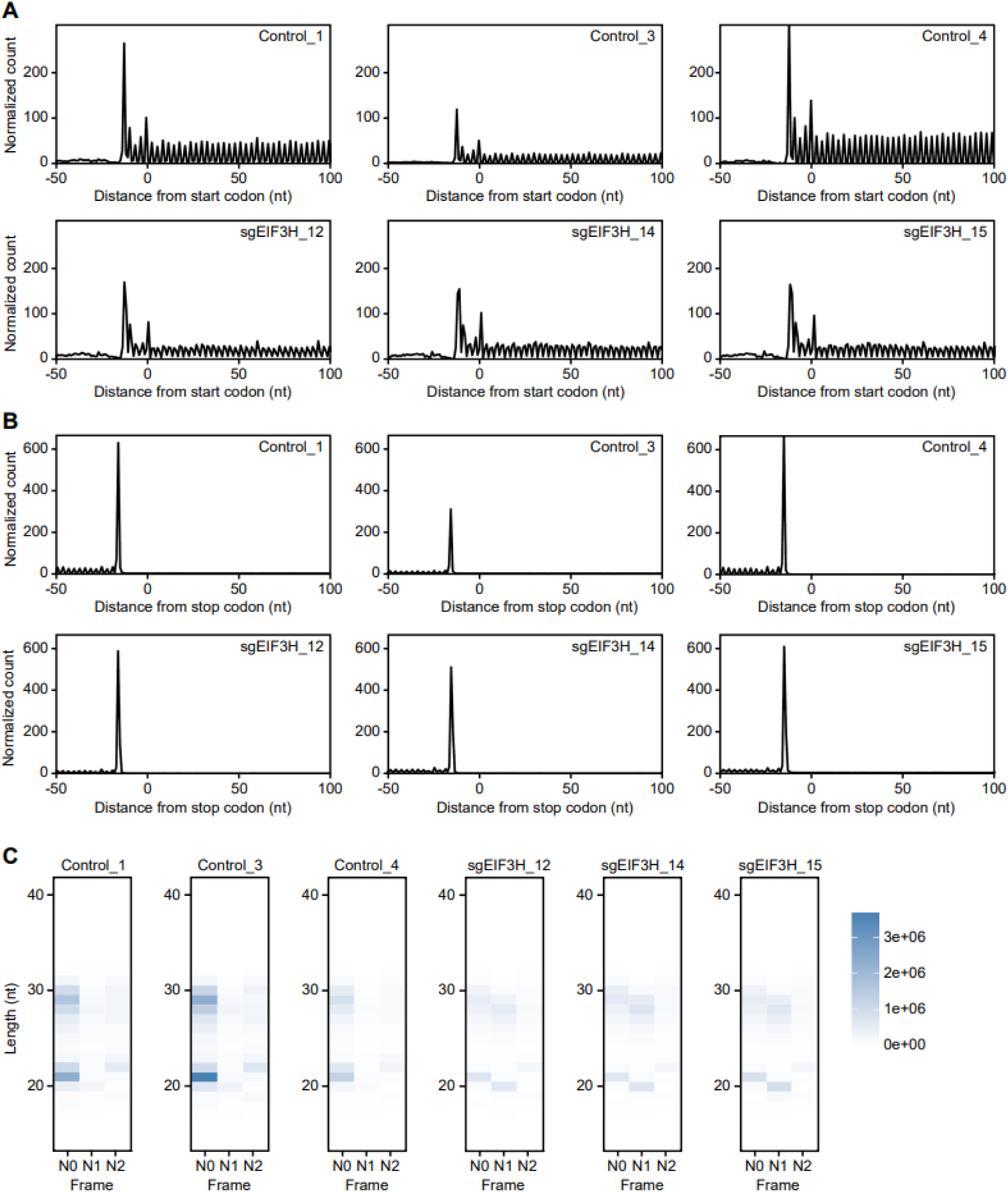
Supporting data for ribosome profiling (Related to Fig. 4) **(A),** Metagene analysis around the start codon. **(B),** Metagene analysis around the stop codon. **(C),** Heatmaps of read counts, categorized by reading frames and read lengths.

**Table S1:** Gene ontology result of upregulated genes in EIF3H KD PSCs on day 3 (related to Fig. 3C)

| Term | Count | Benjamini |
| --- | --- | --- |
| axon guidance | 15 | 5.61E-03 |
| formation of quadruple SL/U4/U5/U6 snRNP | 5 | 5.61E-03 |
| cell differentiation | 38 | 5.61E-03 |
| spliceosomal tri-snRNP complex assembly | 6 | 1.92E-02 |
| positive regulation of bone mineralization | 7 | 1.92E-02 |
| neuron differentiation | 14 | 1.92E-02 |
| cell development | 7 | 2.81E-02 |
| cell adhesion | 25 | 4.36E-02 |

**Table S2:**
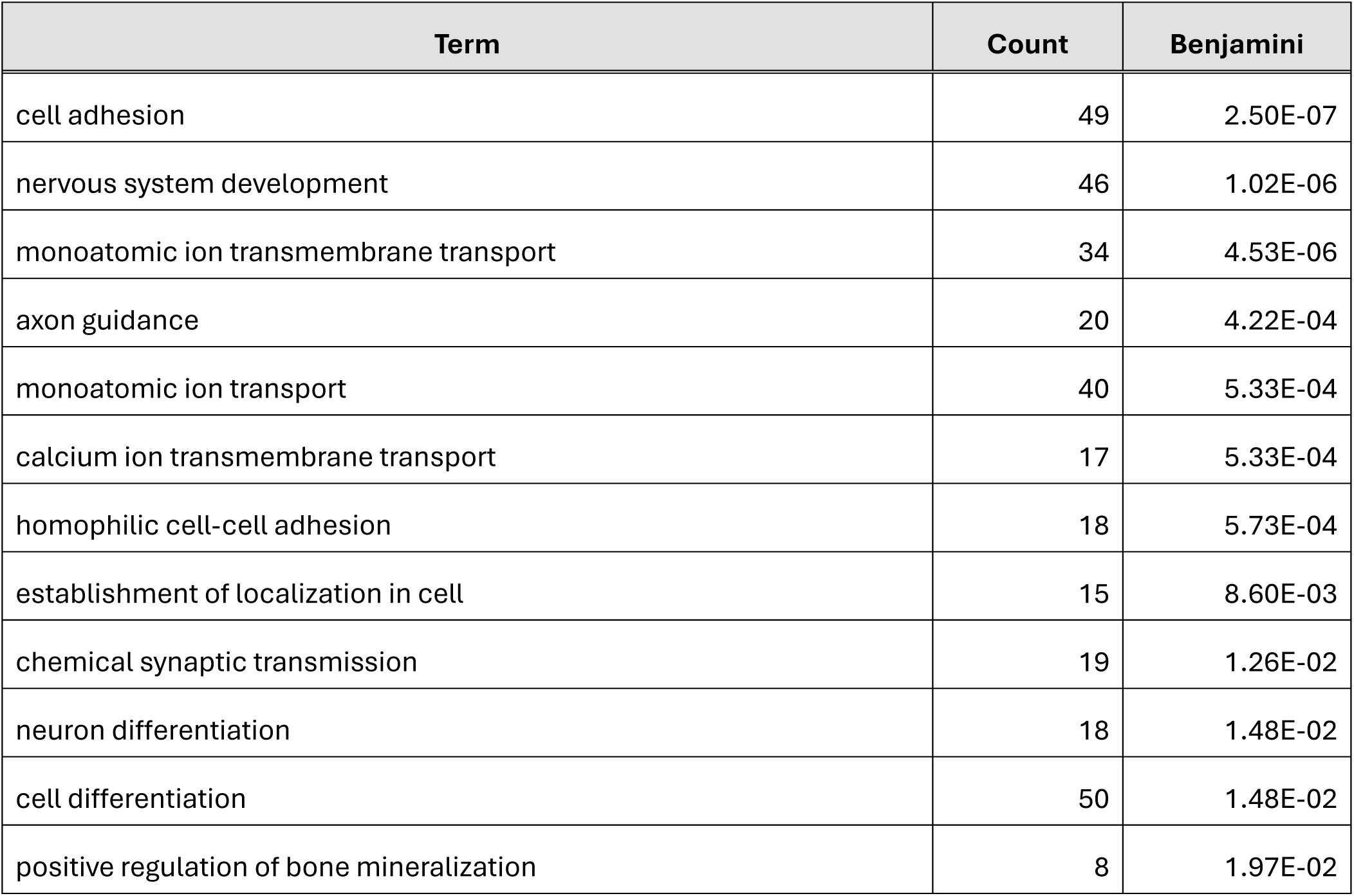
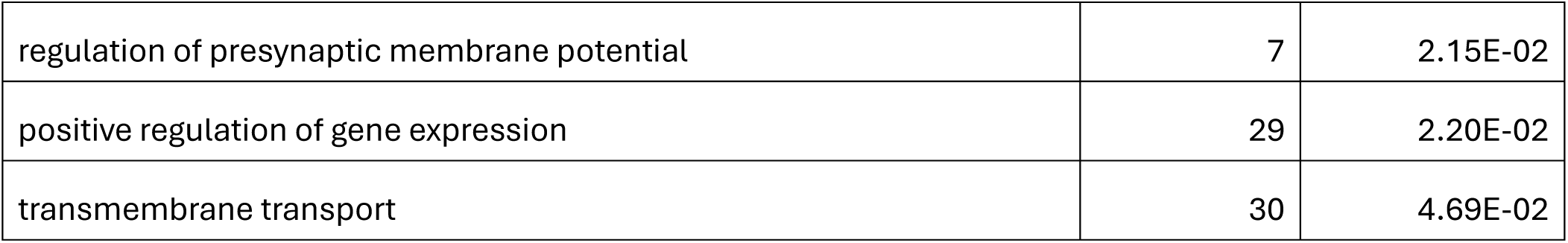
Gene ontology result of upregulated genes in EIF3H KD PSCs on day 5 (related to Fig. 3C)

**Table S3:**
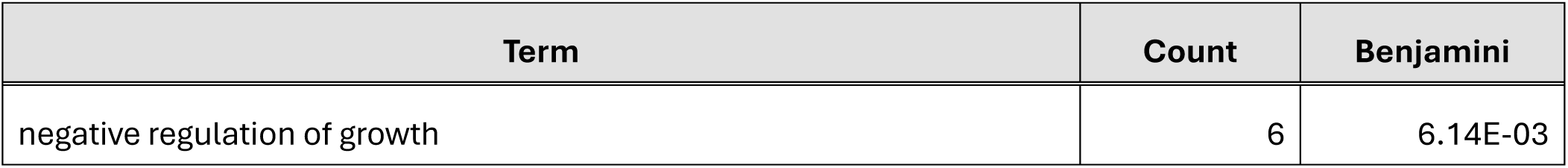
Gene ontology result of downregulated genes in EIF3H KD PSCs on day 5 (related to Fig. 3C)

**Table S4:** Gene ontology result of upregulated genes in EIF3H KD PSCs on day 12 (related to Fig. 3C)

| Term | Count | Benjamini |
| --- | --- | --- |
| nervous system development | 79 | 4.20E-08 |
| angiogenesis | 46 | 2.28E-07 |
| system development | 22 | 6.33E-06 |
| cell adhesion | 73 | 1.44E-05 |
| transmembrane transport | 61 | 2.16E-04 |
| animal organ development | 23 | 2.16E-04 |
| neuron differentiation | 32 | 6.47E-04 |
| monoatomic ion transmembrane transport | 47 | 8.81E-04 |
| negative regulation of cell population proliferation | 48 | 3.51E-03 |
| positive regulation of smooth muscle cell proliferation | 13 | 5.19E-03 |
| central nervous system neuron differentiation | 12 | 5.19E-03 |
| positive regulation of gene expression | 51 | 9.89E-03 |
| bone mineralization | 13 | 1.40E-02 |
| regulation of postsynaptic membrane potential | 13 | 1.55E-02 |
| sensory perception of pain | 11 | 2.23E-02 |
| modulation of chemical synaptic transmission | 19 | 2.23E-02 |
| cell migration | 34 | 2.23E-02 |
| tissue development | 12 | 2.34E-02 |
| cell surface receptor protein tyrosine kinase signaling pathway | 19 | 2.34E-02 |
| axonogenesis | 19 | 2.47E-02 |
| axon guidance | 25 | 2.76E-02 |
| regulation of presynaptic membrane potential | 9 | 3.10E-02 |
| monoatomic ion transport | 59 | 3.37E-02 |
| positive regulation of heart rate | 8 | 4.22E-02 |
| amino acid transport | 13 | 4.84E-02 |

**Table S5:** Gene ontology result of downregulated genes in EIF3H KD PSCs on day 12 (related to Fig. 3C)

| Term | Count | Benjamini |
| --- | --- | --- |
| chemical synaptic transmission | 35 | 4.74E-03 |
| monoatomic ion transmembrane transport | 49 | 4.99E-03 |
| calcium ion transmembrane transport | 25 | 6.21E-03 |
| meiotic cell cycle | 23 | 1.74E-02 |
| monoatomic ion transport | 65 | 3.54E-02 |

**Table S6:** Expression of pluripotency-associated genes in EIF3H KD PSCs on day 12 (related to Fig. 4D)

| Gene | log2FC | Adjusted P-value |
| --- | --- | --- |
| POU5F1 | -0.187345864071276 | 1.84E-02 |
| SOX2 | -0.450684469780254 | 1.24E-06 |
| NANOG | -1.50981546274349 | 4.46E-22 |
| DPPA2 | -2.5260760295195 | 5.98E-33 |
| DPPA4 | -0.83544320613486 | 2.63E-41 |
| IFITM1 | -2.77978775592462 | 1.03E-54 |
| TERT | -0.472349870224465 | 2.27E-02 |
| FGF4 | 4.04491375030254 | 9.82E-50 |
| GABRB3 | 0.0460592366665411 | 6.60E-01 |
| DNMT3B | -0.412395628975791 | 1.23E-09 |
| GDF3 | -1.700616506346 | 4.69E-16 |
| UTF1 | -2.28187439063205 | 1.25E-30 |
| LIN28A | 0.172792374480528 | 1.47E-02 |
| FOXH1 | -0.709298101387206 | 9.84E-18 |
| PRDM14 | -0.277073416221611 | 3.19E-03 |
| L1TD1 | -0.309651317708884 | 6.03E-03 |
| SFRP2 | -0.371339886446705 | 1.92E-03 |
| CD24 | -0.388422835930974 | 1.72E-07 |
| ZIC2 | -0.353394970808224 | 3.45E-02 |

**Table S7:** Gene ontology result of upregulated TE genes in EIF3H KD PSCs (related to Fig. 4D)

| Term | Count | Benjamini |
| --- | --- | --- |
| positive regulation of DNA-templated transcription | 78 | 9.01E-10 |
| chromatin remodeling | 44 | 9.58E-09 |
| negative regulation of transcription by RNA polymerase II | 86 | 1.36E-07 |
| positive regulation of transcription by RNA polymerase II | 92 | 7.84E-06 |
| RNA splicing | 38 | 8.57E-06 |
| regulation of transcription by RNA polymerase II | 110 | 2.86E-05 |
| regulation of DNA-templated transcription | 104 | 2.86E-05 |
| mRNA processing | 41 | 2.06E-04 |
| negative regulation of DNA-templated transcription | 54 | 2.07E-04 |
| chromatin organization | 40 | 2.88E-03 |
| positive regulation of stem cell population maintenance | 11 | 4.58E-03 |
| endocytic recycling | 13 | 4.58E-03 |
| regulation of G1/S transition of mitotic cell cycle | 12 | 4.82E-03 |
| osteoblast differentiation | 18 | 8.58E-03 |
| response to xenobiotic stimulus | 26 | 1.00E-02 |
| proton motive force-driven mitochondrial ATP synthesis | 12 | 1.01E-02 |
| heart development | 27 | 1.14E-02 |
| aerobic respiration | 12 | 1.81E-02 |
| cellular response to epidermal growth factor stimulus | 10 | 2.16E-02 |
| positive regulation of cell differentiation | 13 | 2.57E-02 |
| DNA repair | 35 | 2.57E-02 |
| regulation of apoptotic process | 26 | 4.18E-02 |
| positive regulation of stress fiber assembly | 10 | 4.44E-02 |
| DNA damage response | 42 | 4.44E-02 |
| regulation of mitotic metaphase/anaphase transition | 8 | 4.44E-02 |
| regulation of gene expression | 33 | 4.44E-02 |
| rhythmic process | 17 | 4.44E-02 |
| regulation of G0 to G1 transition | 7 | 4.55E-02 |
| mRNA splicing, via spliceosome | 20 | 4.61E-02 |

**Table S8:** Gene ontology result of downregulated TE genes in EIF3H KD PSCs (related to Fig. 4D)

| Term | Count | Benjamini |
| --- | --- | --- |
| cytoplasmic translation | 56 | 4.01E-39 |
| translation | 69 | 1.06E-26 |
| proton transmembrane transport | 37 | 1.58E-12 |
| proton motive force-driven mitochondrial ATP synthesis | 21 | 3.80E-10 |
| ribosomal small subunit biogenesis | 21 | 1.29E-08 |
| mitochondrial electron transport, NADH to ubiquinone | 14 | 2.67E-05 |
| ribosomal small subunit assembly | 9 | 2.67E-05 |
| aerobic respiration | 16 | 3.50E-05 |
| detoxification of copper ion | 8 | 4.47E-04 |
| cellular respiration | 11 | 1.11E-03 |
| rRNA processing | 22 | 1.24E-03 |
| intracellular zinc ion homeostasis | 10 | 1.77E-03 |
| cellular response to cadmium ion | 8 | 5.18E-03 |
| proton motive force-driven ATP synthesis | 8 | 8.20E-03 |
| cellular response to zinc ion | 8 | 8.20E-03 |
| endoplasmic reticulum to Golgi vesicle-mediated transport | 17 | 1.26E-02 |
| oxidative phosphorylation | 8 | 2.56E-02 |
| 7-methylguanosine cap hypermethylation | 5 | 2.56E-02 |
| mitochondrial electron transport, cytochrome c to oxygen | 7 | 3.56E-02 |
| modification-dependent protein catabolic process | 5 | 4.01E-02 |

**Table S9:** Gene ontology result of downregulated genes in MTF1 KD PSCs (related to Fig. 5B)

| Term | Count | Benjamini |
| --- | --- | --- |
| monoatomic ion transport | 28 | 8.47E-04 |
| cellular response to zinc ion | 7 | 8.47E-04 |
| detoxification of copper ion | 6 | 1.14E-03 |
| negative regulation of growth | 6 | 1.63E-03 |
| intracellular zinc ion homeostasis | 7 | 2.77E-03 |
| cellular response to cadmium ion | 6 | 4.04E-03 |
| monoatomic ion transmembrane transport | 19 | 4.74E-03 |
| cellular response to copper ion | 6 | 5.73E-03 |
| transmembrane transport | 21 | 2.62E-02 |
| sodium ion transport | 10 | 3.58E-02 |

**Table S10:**
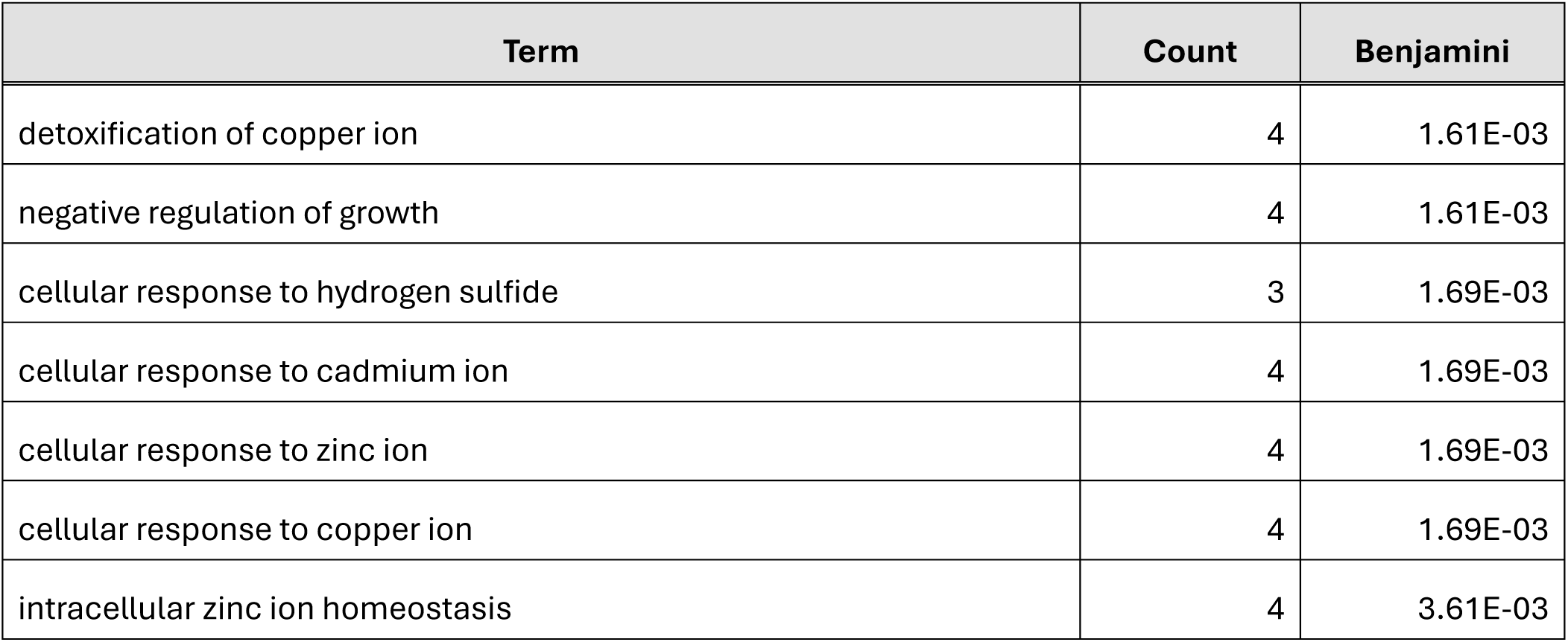
Gene ontology result of downregulated genes common in EIF3H KD and MTF1 KD PSCs on day 5 (related to Fig. 5C)

| Term | Count | Benjamini |
| --- | --- | --- |
| detoxification of copper ion | 4 | 1.61E-03 |
| negative regulation of growth | 4 | 1.61E-03 |
| cellular response to hydrogen sulfide | 3 | 1.69E-03 |
| cellular response to cadmium ion | 4 | 1.69E-03 |
| cellular response to zinc ion | 4 | 1.69E-03 |
| cellular response to copper ion | 4 | 1.69E-03 |
| intracellular zinc ion homeostasis | 4 | 3.61E-03 |

**Table S11:**
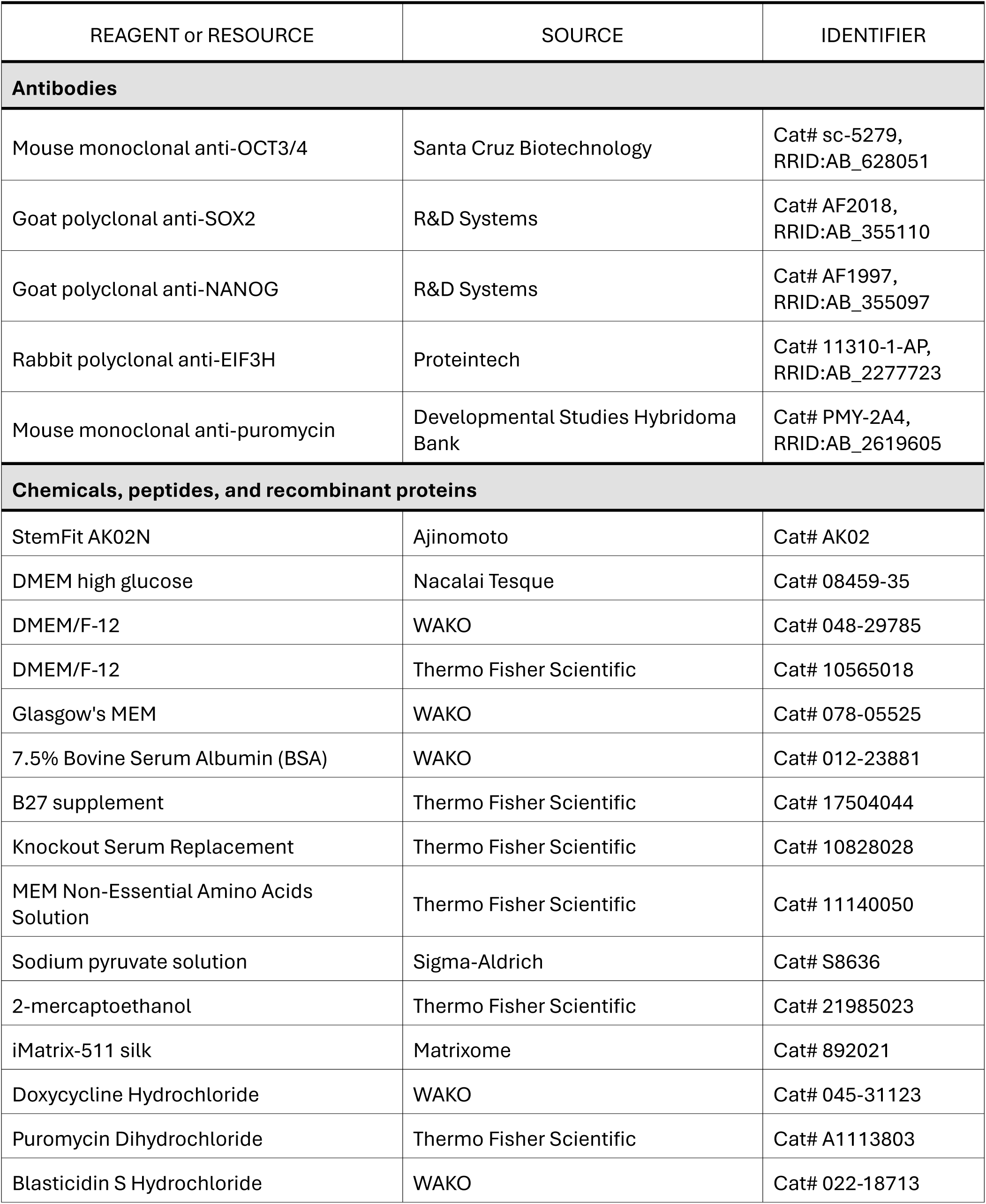
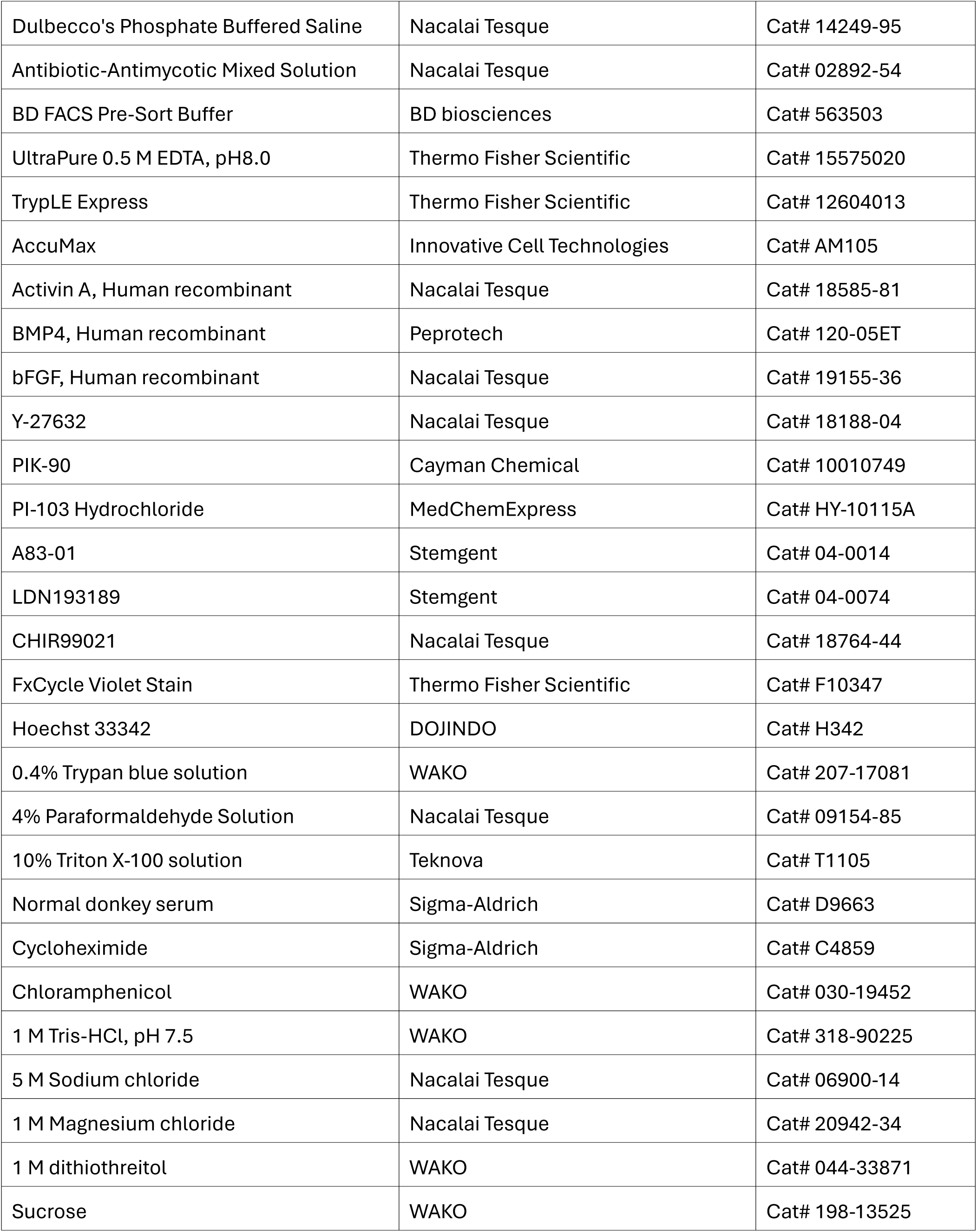
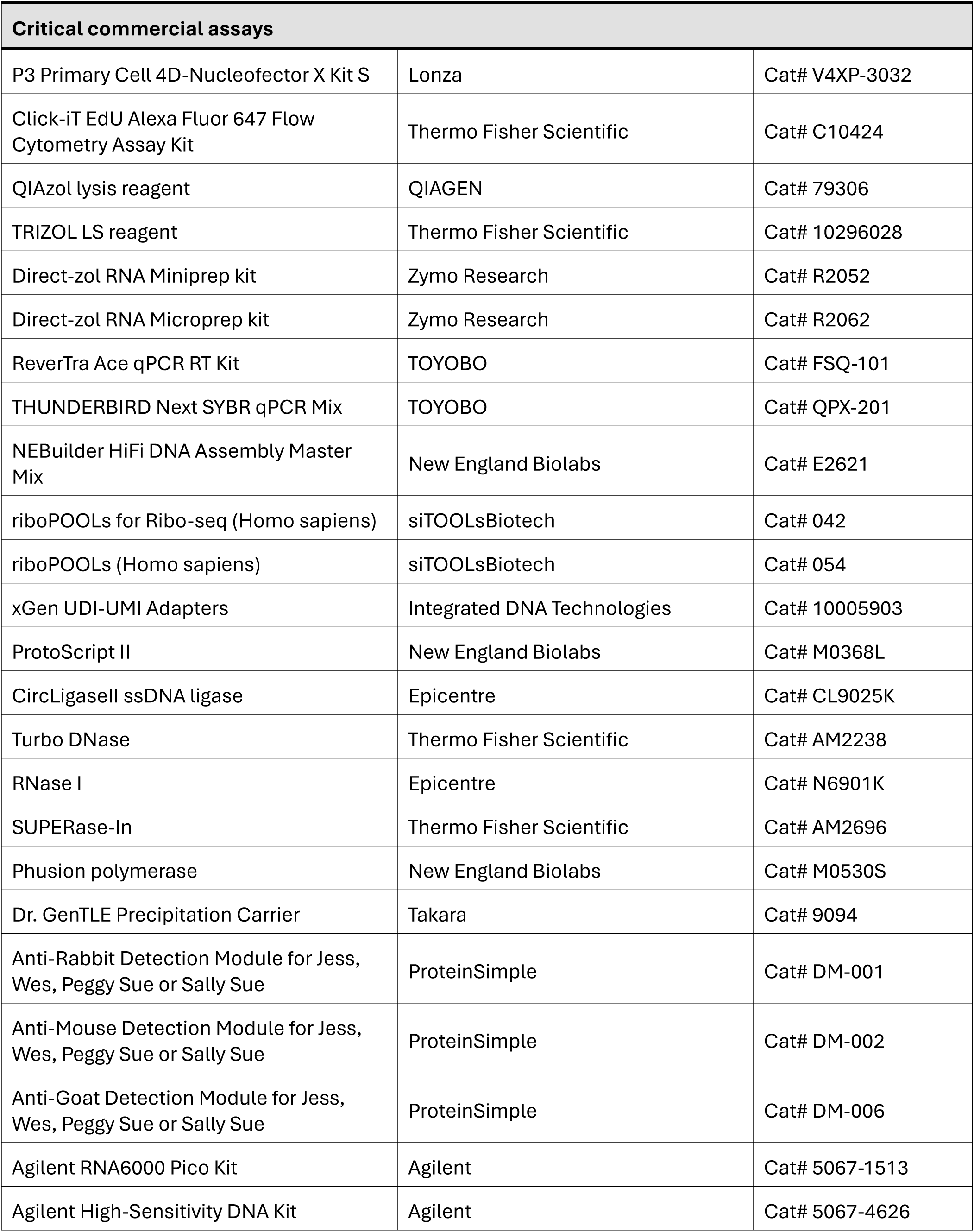
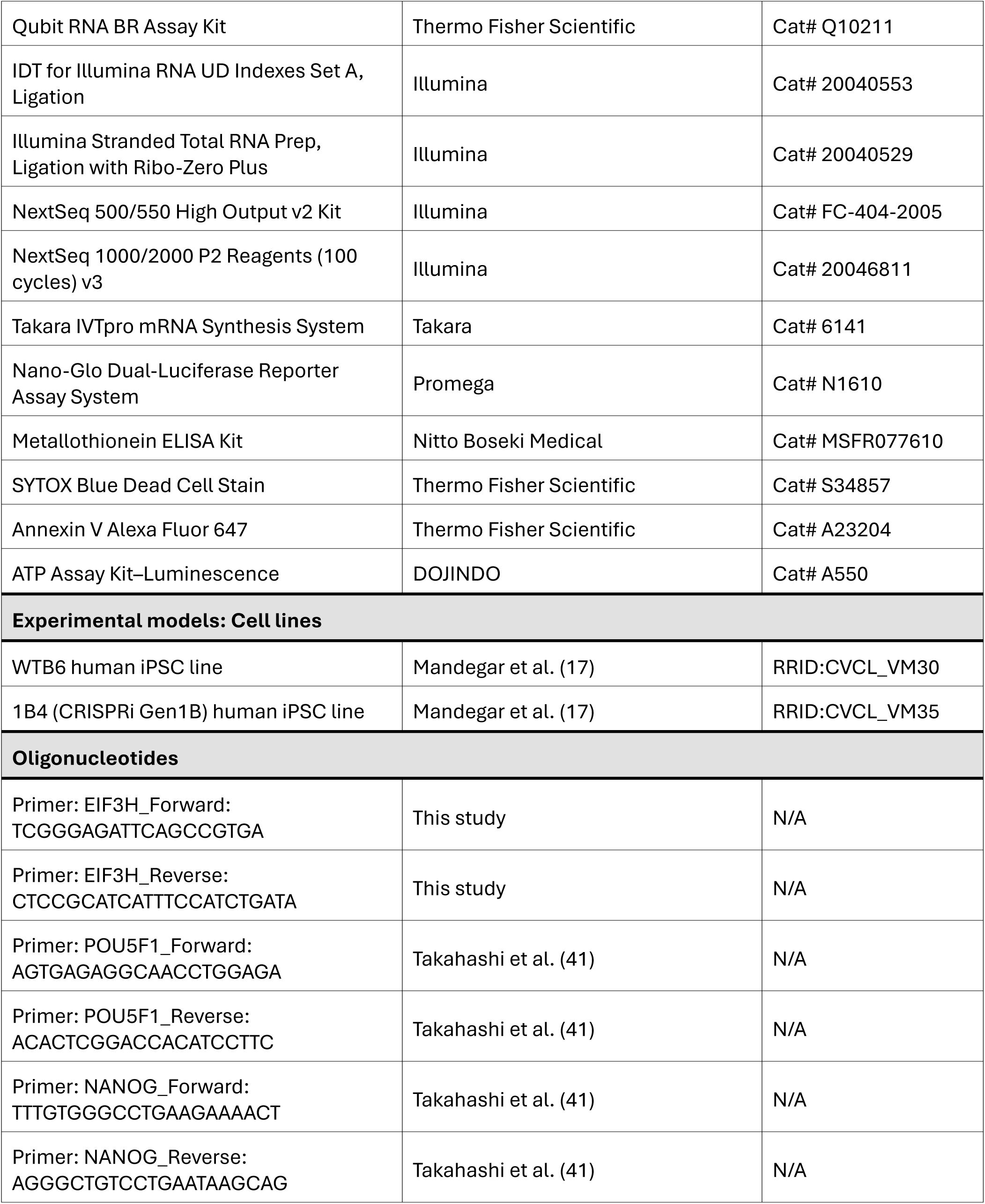
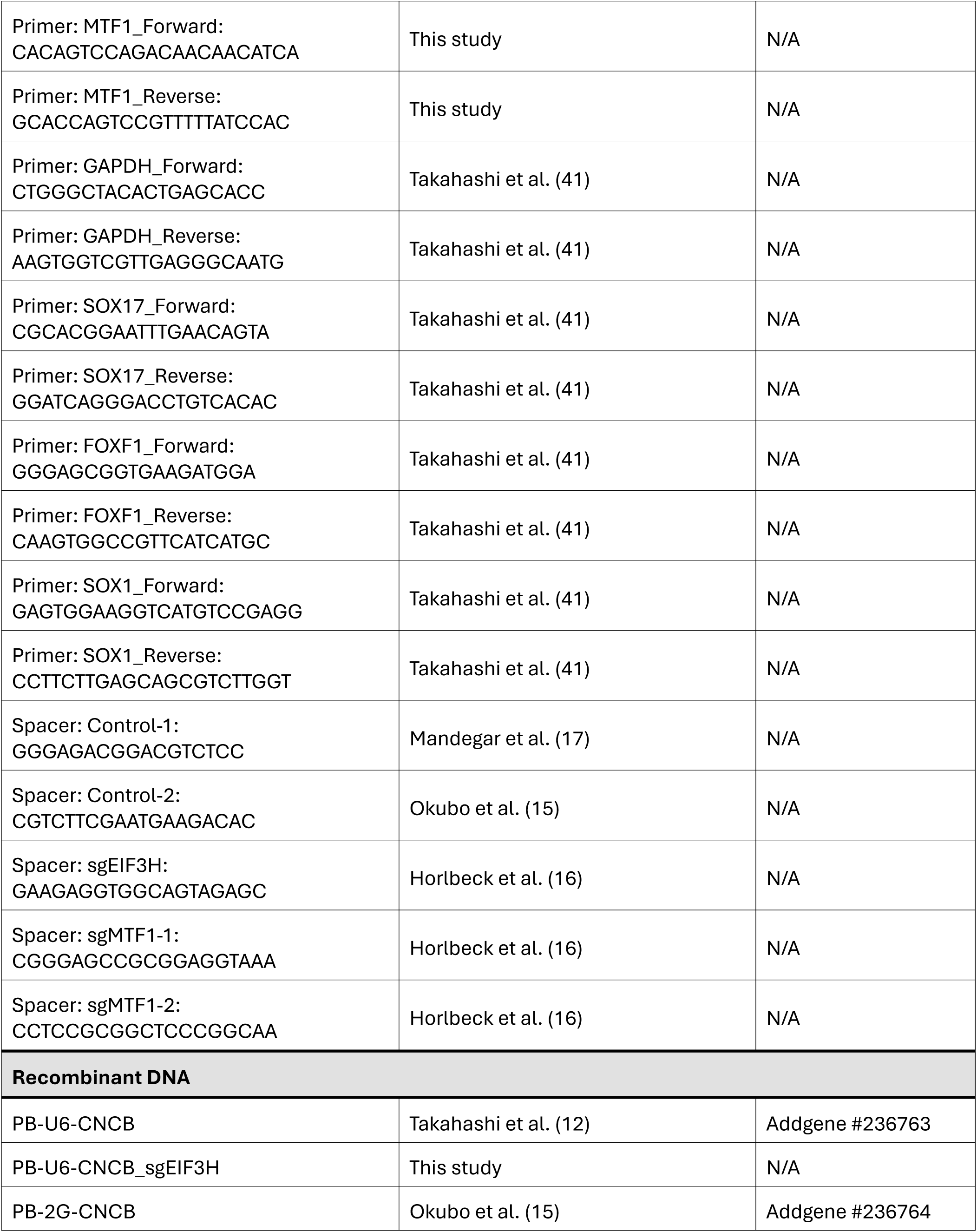
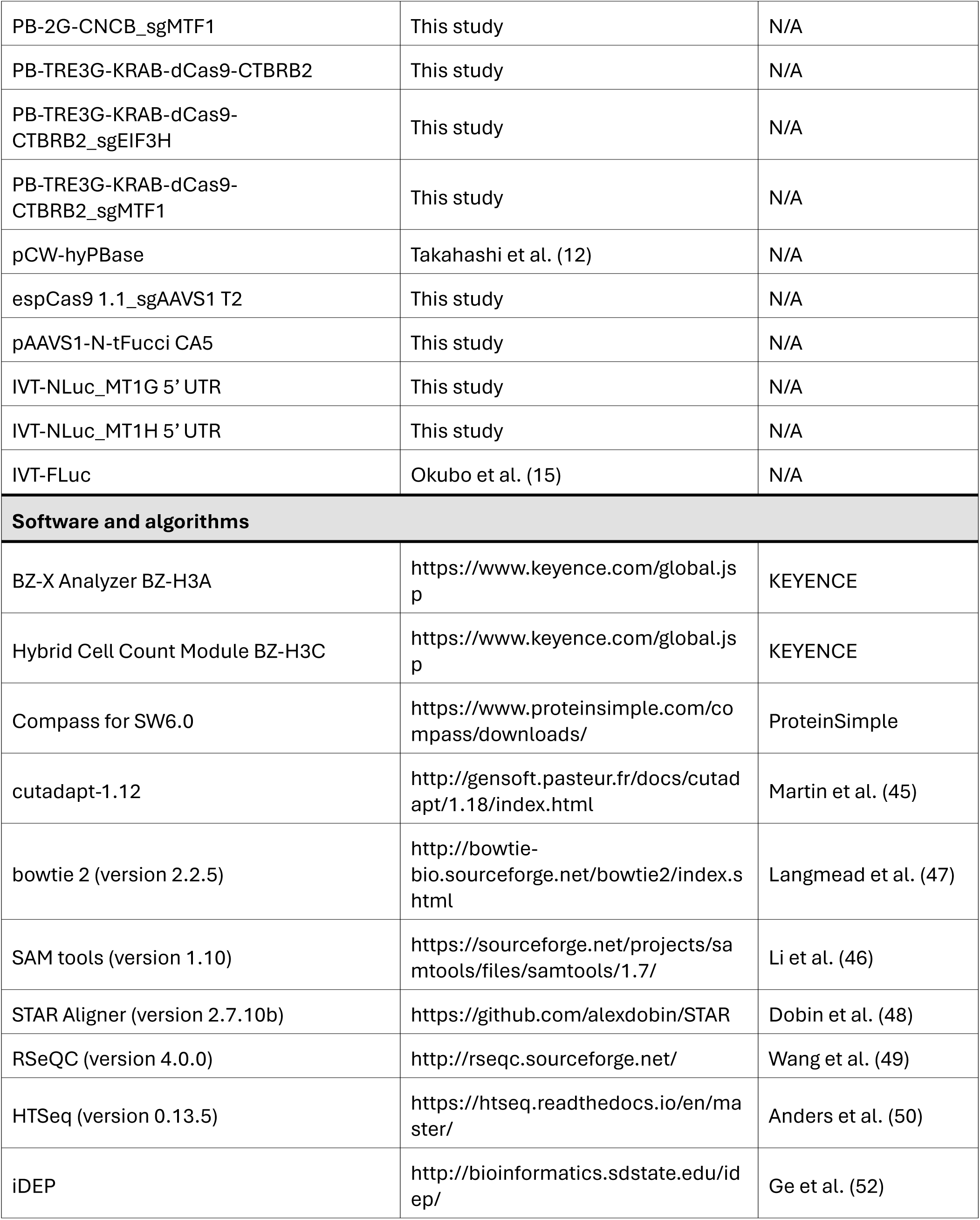
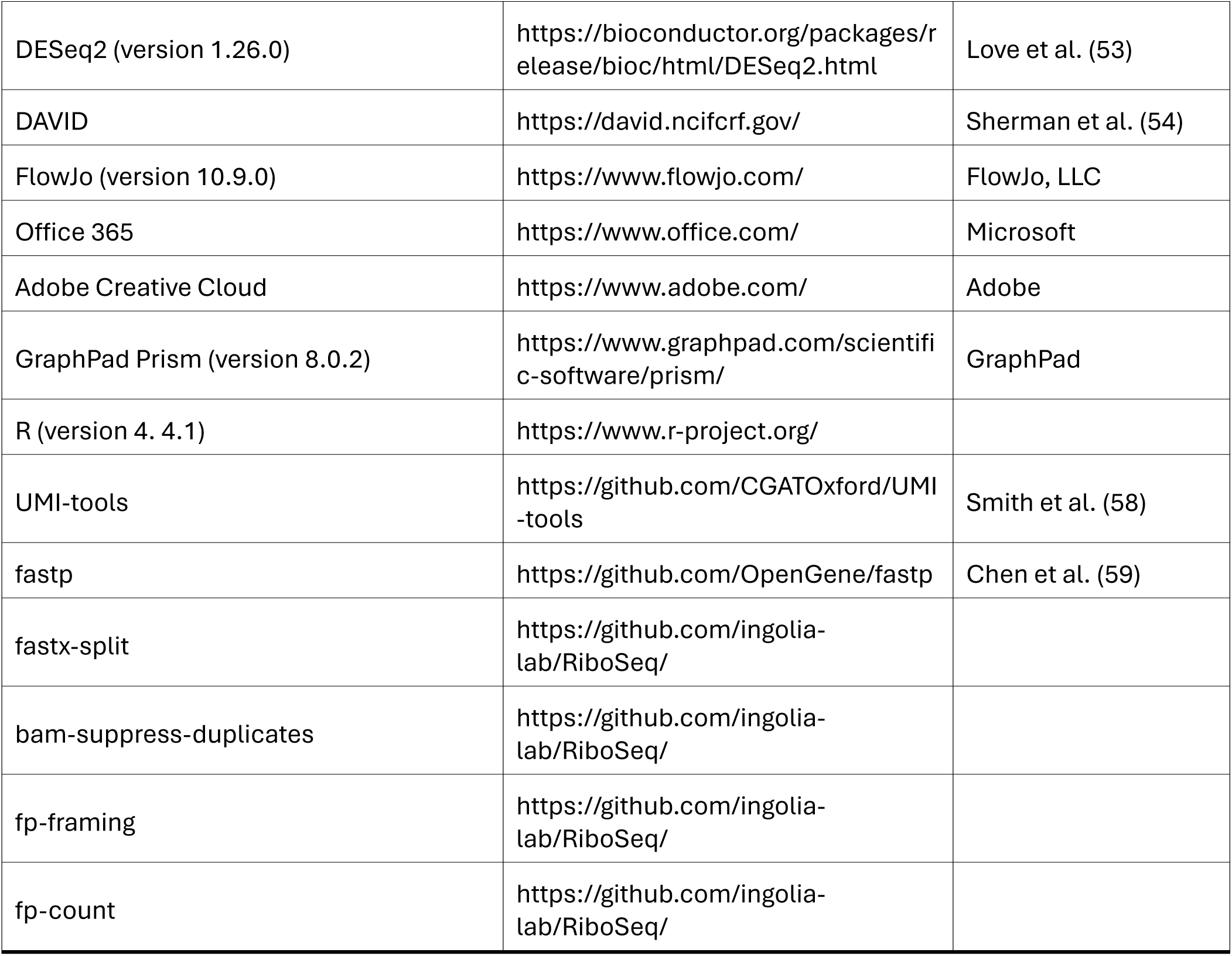
Key resources table.

## METHODS

### Experimental design

To assess the role of translational regulation in primed pluripotent stem cells, we conducted functional analyses targeting EIF3H, which was identified through prior functional screening. Three independent clones of both control and EIF3H KD iPSCs were generated and used in all comparative experiments. For selected assays, the parental 1B4 iPSC line (17) was included as a reference. All experiments were performed with at least 2–3 biological replicates, and the data were analyzed using appropriate statistical methods. No statistical methods were used to predetermine sample size. Experiments were not randomized, but the control and KD samples were processed in parallel to minimize batch effects. The investigators were not blinded to group allocation during the experiments. For comparisons between two groups, two-tailed unpaired Student’s t-tests were used unless otherwise stated. For multiple comparisons, one-way analysis of variance followed by Tukey’s post-hoc test was applied. P-values < 0.05 were considered statistically significant. The exact *n* values, statistical tests, and P-values are indicated in the figure legends.

### Cell lines

Human iPSC lines WTB6 and 1B4 (a CRISPRi Gen1 derivative of WTB6) were used in this study. To establish inducible CRISPRi iPSC lines targeting the gene of interest, 1B4 human iPSCs (passage 23) were transfected with a vector encoding a U6 promoter-driven sgRNA, a CAG promoter-driven fluorescent reporter, and a selectable marker using a PiggyBac-based delivery system as previously described (17). Selection of the appropriate drug commenced two days after transfection and continued until all non-transfected cells were eliminated. Single-cell-derived colonies exhibiting uniform fluorescence were subsequently picked and expanded. Knockdown of the target gene was induced by treating cells with 1 μg/mL Dox (WAKO) for the indicated time periods. Clones within 20 passages following subcloning were used for experiments. sgRNA spacer sequences used in this study are listed in Table S11.

### Cell culture

Human PSCs were maintained in StemFit AK02N medium (Takara Bio) under standard culture conditions as previously described (41, 42). The cells were maintained at 37°C in a humidified incubator with 5% CO₂ and ambient O₂. The medium was changed every day. For passaging, the cells were washed with Dulbecco’s phosphate-buffered saline (D-PBS; Nacalai Tesque), dissociated with TrypLE Express (Thermo Fisher Scientific) for 10 min at 37°C, and resuspended in DMEM/F-12 (Nacalai Tesque) supplemented with 0.1% bovine serum albumin (BSA; Wako). After centrifugation, cells were replated onto iMatrix-511 silk-coated plates (1.67 µg/mL; Matrixome) in StemFit AK02N medium supplemented with 10 µM Y-27632 (Nacalai Tesque). Y-27632 was added 24 h after passage.

### Generation of tFucci reporter iPSC line

A fragment of tFucci (CA)5 (courtesy of Atsushi Miyawaki; Addgene #153521) (19), followed by a woodchuck hepatitis virus post-transcriptional regulatory element (WPRE) fragment, was subcloned into the EcoRI sites of pAAVS1-Nst-CAG-DEST (Addgene #80489) using NEBuilder HiFi DNA Assembly technology (New England Biolabs). The plasmid, along with the eSpCas9 (1.1) plasmid (courtesy of Feng Zhang; Addgene #71814) encoding AAVS1-T2 sgRNA, was transfected into WTB6 iPSCs (P59). Three days after transfection, the cells were treated with 250 μg/mL G418 (Wako) until the non-transfected cells were completely eliminated. Subsequently, single-cell-derived colonies were isolated and expanded. The clones were evaluated by genotyping and karyotyping, and the copy number of the transgenes was confirmed using digital droplet PCR.

### Cell division time measurement

Cell cycle duration was measured using the tFucci reporter iPSC line. To induce EIF3H KD, cells were treated with Dox for the indicated number of days prior to imaging. Live-cell imaging was performed using a BioPipeline LIVE imaging system (Nikon), with cells maintained at 37°C under 5% CO₂.

Phase-contrast and fluorescence images were acquired every 15 min for 48 h using a 10× objective lens. For each biological replicate, individual cells that were spatially isolated and that remained within the imaging field throughout the observation period were selected for analysis. Cell division time was defined as the interval between two consecutive cytokinesis events, as determined by morphological changes in phase-contrast images and the corresponding transitions in tFucci fluorescence signals.

For each cell line, at least three independent cells per biological replicate were manually tracked using the NIS Viewer (Nikon).

### Growth competition assay

A 1:1 mixture of 1B4 cells (wild-type, GFP-negative) and control or KD cells (GFP-positive) was seeded into a single well of a six-well plate at a density of 4 × 10^5^ cells. The mixed population was cultured in a medium containing Dox and passaged every three days under the same conditions. Cells were continuously maintained until day 12. At each passage, the proportion of GFP-positive cells in the collected cell population was quantified by flow cytometry.

### Endoderm differentiation

Endoderm differentiation was performed as previously reported with slight modifications (36, 37). Primed PSCs were plated at a density of 1 × 10^6^ cells per well on iMatrix 511-coated 6-well plates in StemFit AK02N medium supplemented with 10 μM Y-27632. On the following day, cells were washed once with DMEM/F-12, and the medium was replaced with Differentiation Medium 1 (DM1) comprising DMEM/F-12 (Thermo Fisher Scientific), 2% B27 supplement (Thermo Fisher Scientific), 1% MEM Non-Essential Amino Acids (NEAA, Thermo Fisher Scientific), and 0.1 mM 2-mercaptoethanol (2-ME, Thermo Fisher Scientific), supplemented with 100 ng/mL Activin A (Nacalai Tesque), 3 μM CHIR99021 (Nacalai Tesque), 20 ng/mL bFGF (Nacalai Tesque), and 50 nM PI-103 (Cayman Chemical). After 24 h, the cells were washed with DMEM/F-12, and DM1 containing 100 ng/mL Activin A and 250 nM LDN193189 (Stemgent) was added. Two days later, after further washing with DMEM/F-12, the cells were maintained in DM1 supplemented with 100 ng/mL Activin A for an additional 48 h.

### Mesoderm differentiation

Mesodermal differentiation was performed following previously described protocols, with minor modifications (37, 38). Primed PSCs were plated one day before induction at a density of 1 × 10^6^ cells per well on iMatrix 511-coated 6-well plates in StemFit AK02N medium supplemented with 10 μM Y-27632. On the following day, cells were washed once with DMEM/F-12 and cultured in DM1 medium containing 30 ng/mL Activin A, 40 ng/mL BMP4 (PeproTech), 6 μM CHIR99021, 20 ng/mL bFGF, and 100 nM PIK-90 (MedChemExpress) for 24 h. After this period, cells were washed with DMEM/F-12 and maintained in DM1 supplemented with 40 ng/mL BMP4, 1 μM A83-01, and 4 μM CHIR99021 for an additional 48 h. After another wash with DMEM/F-12, the cells were cultured in DM1 containing 40 ng/mL BMP4 for 48 h.

### Ectoderm differentiation

Neuroectoderm differentiation was performed as previously described with minor modifications (39, 40). One day before induction, primed PSCs were seeded at a density of 1 × 10^6^ cells per well on iMatrix 511-coated 6-well plates in StemFit AK02N medium supplemented with 10 μM Y-27632. On the following day, cells were washed once with DMEM/F-12 and then cultured in Glasgow’s MEM (Wako) supplemented with 8% Knockout Serum Replacement (KSR; Thermo Fisher Scientific), 1 mM sodium pyruvate (Sigma-Aldrich), 1% MEM nonessential amino acids, 0.1 mM 2-mercaptoethanol, 1 μM A83-01, and 250 nM LDN193189. The differentiation medium was refreshed daily for a total of 5 days.

### RNA extraction and quantitative PCR

Cells were rinsed once with D-PBS and subsequently lysed in QIAzol reagent (QIAGEN). Total RNA was purified using the Direct-zol RNA Miniprep Kit (Zymo Research), including on-column DNase digestion, according to the manufacturer’s protocol. For reverse transcription, 1 μg of total RNA was used with ReverTra Ace qPCR RT Master Mix (TOYOBO). Quantitative PCR was performed using gene-specific primers listed in Table S11 and THUNDERBIRD Next SYBR qPCR Mix (TOYOBO) on a QuantStudio 5 Real-Time PCR System (Applied Biosystems).

Cycle threshold (Ct) values were normalized to *GAPDH* as endogenous controls, and relative gene expression levels were calculated using the ΔΔCt method. Fold changes are expressed relative to the appropriate control samples. Each qPCR reaction was performed in technical duplicates, and melting curve analysis confirmed single-product amplification.

### Size-based protein analysis

Cells were rinsed once with D-PBS and lysed in radioimmunoprecipitation assay (RIPA) buffer (Sigma-Aldrich) supplemented with a protease inhibitor cocktail (Sigma-Aldrich). The lysates were clarified by centrifugation at 15,300 ×*g* for 15 min at 4°C, after which the supernatant was transferred to a fresh microtube. Protein concentrations were determined using the Pierce BCA Protein Assay Kit (Thermo Fisher Scientific) and quantified on an EnVision 2104 plate reader (Perkin Elmer) following procedures adapted from previously established protocols.

For the quantitative detection of specific proteins, samples were processed using either the Wes or Jess automated capillary electrophoresis platform (ProteinSimple) equipped with 12–230 kDa separation modules. A total of 2 μg of lysate was loaded per capillary, together with the following primary antibodies listed in Table S11: rabbit polyclonal anti-EIF3H (1:250, Proteintech), mouse monoclonal anti-puromycin (1:20, Developmental Studies Hybridoma Bank), mouse monoclonal anti-OCT3/4 (1:500, Santa Cruz Biotechnology), goat polyclonal anti-SOX2 (1:40, R&D Systems), goat polyclonal anti-NANOG (1:40, R&D Systems), and rabbit monoclonal anti-VINCULIN (1:250, Cell Signaling Technology). Signal acquisition and quantitative analysis were performed using Compass for SW6.0 software (ProteinSimple).

### Apoptosis assay

The cells were washed with D-PBS, collected by centrifugation, and stained using the Annexin V Conjugates for Apoptosis Detection Kit (Thermo Fisher Scientific) following the manufacturer’s instructions. The samples were analyzed by flow cytometry, and the data were processed using FlowJo software.

### Cell cycle analysis

Cell cycle profiling was performed using the Click-iT EdU Alexa Fluor 647 Flow Cytometry Assay Kit (Thermo Fisher Scientific) following a protocol modified from previously reported methods (15). Cells were plated in 6-well plates at a density of 5 × 10^5^ cells per well and maintained for 5 days in StemFit AK02N medium supplemented with doxycycline. To label S-phase cells, cultures were exposed to 10 µM 5-ethynyl-2′-deoxyuridine (EdU) for 135 min at 37°C. After incorporation, the cells were collected, washed with 1% BSA in D-PBS, and pelleted by centrifugation. Cell pellets were fixed for 15 min at room temperature using the Click-iT fixative solution and rinsed again with 1% BSA. Permeabilization was achieved by incubating the samples with 1× Click-iT Perm/Wash reagent for 15 min at room temperature. EdU detection was performed using a Click-iT reaction cocktail containing a Copper Protectant, Alexa Fluor 647 picolyl azide, and the Click-iT buffer additive. Following the reaction, cells were washed with 1× Click-iT Perm/Wash reagent, stained with 1 μg/mL FxCycle Violet (Thermo Fisher Scientific) for 30 min at room temperature to determine DNA content, and subsequently analyzed by flow cytometry. A total of 1 × 10^4^ events were acquired on a FACS Aria II (BD Biosciences) using BD FACSDiva software. Fluorescence derived from EdU-Alexa Fluor 647 and FxCycle Violet was detected using APC and Pacific Blue channels, respectively. Data processing and gating strategies were performed using FlowJo software.

### Messenger RNA synthesis and luciferase assay

For the luciferase reporter assay, a T7 promoter–driven NanoLuc luciferase (NLuc) construct was generated. The 5′ UTR sequences of human MT1G (72 bp from NM_001301267.2) and MT1H (71 bp from NM_005951.2) were inserted between the T7 promoter and NLuc using NEBuilder assembly cloning (New England Biolabs), following the manufacturer’s instructions. To prepare a normalization control plasmid, a firefly luciferase (FLuc) gene was introduced into the linearized vector of the IVTpro mRNA Synthesis System (Takara). Messenger RNAs were synthesized using these plasmids according to the manufacturer’s protocol, incorporating CleanCap Reagent AG (Trilink BioTechnologies) as the cap analog and N1-methylpseudouridine 5’-triphosphate sodium salt (TCI Chemicals) in place of uridine triphosphate. The integrity and quality of the synthesized mRNAs were confirmed using a bioanalyzer (Agilent Technologies).

After 2 days of Dox treatment, control and sgEIF3H PSCs were replated onto iMatrix 511–coated 24-well plates at a density of 7.5 × 10^4^ cells per well in StemFit AK02N medium supplemented with 10 μM Y-27632 and Dox. The following day (day 3 post-Dox induction), the cells were transfected with an mRNA mixture consisting of 125 ng each of the NLuc reporter and FLuc control mRNAs using Lipofectamine Stem Reagent (Thermo Fisher Scientific). Transfection was performed using 1 μL Lipofectamine Stem Reagent per well (24-well plate), and complexes were incubated for 10 min before addition to cells. After an additional 24 h, the cells were lysed in 1× Passive Lysis Buffer (Promega), and luciferase activity was quantified using a GloMax Discover Microplate Reader (Promega) and the Nano-Glo Dual-Luciferase Reporter Assay System (Promega).

### iATP mRNA transfection and flow cytometry

We synthesized mRNA encoding (mito).iATPSnFR2.A95A.A119L.miRFP670nano3, a mitochondria-targeted, low-affinity, ratiometric ATP sensor (27) using the IVTpro mRNA Synthesis System. We introduced 2 μg of mRNA into the cells growing in 24-well plate using Lipofectamine Stem Reagent (Thermo Fisher Scientific) according to the manufacturer’s instruction. On the following day, flow cytometry was performed using a BD LSR Fortessa flow cytometer (BD Biosciences). Fluorescence signals were acquired in the FITC and APC channels, and the median fluorescence intensity for each channel was quantified.

### ATP measurement

Intracellular ATP was quantified using the ATP Assay Kit–Luminescence (DOJINDO), following the manufacturer’s instructions. The working solution was prepared by reconstituting the substrate with assay buffer and adding an enzyme solution.

ATP standards (0–2.5 µmol/L) were generated from a 1 mmol/L ATP stock by serial dilution in serum-free medium. Cells were assayed directly in white 96-well plates; 100 µL of sample or standard was mixed with 100 µL of working solution in each well. Plates were shaken for 2 min and incubated for 10 min at 25°C, and luminescence was measured using an Envision microplate reader (PerkinElmer). ATP concentrations were calculated from the standard curve and expressed as mean ± SD from triplicate measurements. ATP values were not further normalized; cells were seeded at equal densities under all conditions before measurement.

### Puromycin incorporation

For puromycin incorporation analysis, cells cultured in three separate wells were rinsed twice with pre-warmed D-PBS. One well was then incubated in StemFit AK02N medium supplemented with 100 μg/mL cycloheximide (CHX; Sigma-Aldrich), while the remaining two wells received medium without additives. Following a 10-min incubation at 37°C, puromycin (1 μM; Thermo Fisher Scientific) was added to the CHX-treated culture and to one of the untreated wells, and incubation proceeded for an additional 30 min at 37°C. At the end of the treatment period, the cells were washed with ice-cold D-PBS and lysed in RIPA buffer containing a protease inhibitor cocktail. Proteins were subsequently analyzed using size-based separation as described above.

## MT ELISA

Cells were dissociated using TrypLE Express (Thermo Fisher Scientific), washed once with PBS, and collected by centrifugation. The cell pellets were then resuspended in lysis buffer consisting of 50 mM Tris-HCl (pH 7.4), 150 mM NaCl, 1% NP-40, 1 mM EDTA, and 1× protease inhibitor cocktail (Complete EDTA-free, Roche). The lysates were then clarified by centrifugation at 15,000 ×*g* for 15 min at 4°C. The resulting supernatants were collected and stored at −80°C until use.

Total protein concentrations were determined using a Qubit Protein Broad Range (BR) Assay Kit (Thermo Fisher Scientific), following the manufacturer’s instructions. Equal amounts of protein samples were used to measure MT levels using a Metallothionein ELISA Kit (Nitto Boseki Medical), following the manufacturer’s protocol. The absorbance was measured at 450 nm using a GloMax Discover microplate reader (Promega).

### RNA sequencing

Cells were lysed in QIAzol reagent, and total RNA was purified as described above. RNA quality was assessed using the Agilent RNA 6000 Pico Kit on a Bioanalyzer 2100 (Agilent Technologies). Library preparation and downstream analyses were performed as previously described (43, 44). Briefly, 100 ng of DNase-treated total RNA was used for library construction using the Illumina Stranded Total RNA Prep Ligation with the Ribo-Zero Plus kit (Illumina). Library quality was evaluated using an Agilent High-Sensitivity DNA Kit (Agilent Technologies), and sequencing was performed on a NextSeq 500/550 High Output v2 system (Illumina), NextSeq 1000/2000 using P2 Reagents (100 cycles) v3 (Illumina), or a HiSeq X platform (Illumina).

Adapter sequences were trimmed using Cutadapt (45). Reads mapping to ribosomal RNA were removed using SAMtools (46) and Bowtie2 (47). The remaining reads were aligned to the human genome (hg38) using STAR (48). Quality control was performed using RSeQC (49). Gene-level counts were obtained using HTSeq (50) with GENCODE annotation (51).

Downstream analyses, including normalization, principal component analysis, heatmap visualization, and hierarchical clustering, were performed using iDEP. Differential expression analysis was performed using the DESeq2 algorithm implemented in iDEP (52). GO enrichment analysis was performed using DAVID.

For days 3 and 5 RNA-seq analyses, time-matched control datasets from our previous study (15) were used as the control group for comparison.

### Polysome profiling

Polysome fractionation was performed using a protocol adapted from previously published methods with minor modifications (56). For each condition (control or sgEIF3H PSCs), one semiconfluent 100-mm culture dish was transferred onto a CoolBox XT Workstation (Biocision) maintained at 4°C. Cells were rinsed gently with 5 mL of ice-cold D-PBS and then detached by scraping in 0.6 mL of ice-cold lysis buffer containing 20 mM Tris-HCl (pH 7.5), 150 mM NaCl, 5 mM MgCl₂ (Nacalai Tesque), 1 mM dithiothreitol (Wako), 1% Triton X-100, a protease inhibitor mixture, 100 μg/mL cycloheximide, and 100 μg/mL chloramphenicol. The suspension was transferred into pre-chilled 1.5-mL DNA LoBind tubes (Eppendorf). Lysates were incubated for 15 min on ice with Turbo DNase (25 U/mL; Thermo Fisher Scientific) and clarified by centrifugation at 20,000 ×*g* for 10 min at 4°C. Supernatants were immediately collected, flash-frozen in liquid nitrogen, and stored at −80°C until use. Continuous 10–45% sucrose gradients were prepared in 14 × 95-mm polyclear tubes (Seton) by combining 10% and 45% sucrose solutions (Sigma-Aldrich) in polysome buffer (25 mM Tris-HCl, pH 7.5, 150 mM NaCl, 15 mM MgCl₂) supplemented with 1 mM DTT and 100 μg/mL cycloheximide, using a Biocomp Gradient Master. After thawing, the RNA concentrations of the lysates were quantified using the Qubit RNA BR Assay Kit (Thermo Fisher Scientific). Equal RNA input was ensured by loading 40 μg of RNA (in 300 μL) onto each gradient. Samples were centrifuged in a P40ST rotor (himac) at 36,000 rpm for 2.5 h at 4°C. The gradients were fractionated using a Biocomp Piston Gradient Fractionator, and the absorbance profiles at 254 nm were used to visualize the free ribosomal subunits, monosomes, and polysomes. The area under each curve was quantified using GraphPad Prism 8.0.2.

### Ribosome profiling

Ribosome profiling was performed as previously described (56, 57). Cells were lysed in a buffer containing 20 mM Tris-HCl (pH 7.5), 150 mM NaCl, 5 mM MgCl₂, 1 mM DTT, 1% Triton X-100, 100 μg/mL chloramphenicol, and 100 μg/mL CHX, followed by a 15-min DNase treatment on ice. RNA concentrations in the lysates were measured using the Qubit RNA BR Assay Kit (Thermo Fisher Scientific). Ten micrograms of RNA were digested with RNase I (Epicentre) for 45 min at 25°C. Ribosome-protected fragments (RPFs) were concentrated by ultracentrifugation through a sucrose cushion [20 mM Tris-HCl (pH 7.5), 150 mM NaCl, 5 mM MgCl₂, 1 mM DTT, 20 U/mL SUPERase-In (Thermo Fisher Scientific), 1 M sucrose, 100 μg/mL chloramphenicol, and 100 μg/mL CHX]. Pellets were resuspended in pellet buffer [20 mM Tris-HCl (pH 7.5), 300 mM NaCl, 5 mM MgCl₂, 1 mM DTT, 1% Triton X-100, and 20 U/mL SUPERase-In] and purified using the Direct-zol RNA Microprep Kit (Zymo Research). Purified RNAs were separated by electrophoresis, and fragments of 17–34 nt were excised and purified using Dr. GenTLE Precipitation Carrier (Takara). RPFs were ligated to linker oligonucleotides containing an inner index and a unique molecular identifier (UMI). rRNA was depleted using riboPOOLs for Ribo-seq (siTOOLs Biotech). The remaining RNAs were reverse-transcribed using ProtoScript II (New England Biolabs) and circularized with circLigase2 (Epicentre). cDNA templates were amplified using Phusion polymerase (New England Biolabs) with indexed primers.

### TE Calculation

To calculate TE, parallel RNA-seq libraries were prepared from RNA extracted from the lysis buffer using TRIzol LS Reagent (Thermo Fisher Scientific) and the Direct-zol RNA Microprep Kit. RNA-seq libraries were prepared according to the manufacturer’s instructions, except that rRNA depletion was performed using riboPOOLs for RNA-seq (siTOOLs Biotech) and adapter ligation was performed with xGen UDI-UMI Adapters (Integrated DNA Technologies). Sequencing was performed using an RNA-seq workflow.

For RNA-seq libraries, adapter trimming was performed using Cutadapt. Reads mapping to rRNA were removed using Bowtie2 and SAMtools. The remaining reads were aligned to the human genome (hg38) using STAR, and PCR duplicates were removed using UMI information with UMI-tools (58).

For ribosome profiling libraries, reads were demultiplexed using the inner index, and adapter trimming was performed using fastp (59). Reads mapping to rRNA were removed using Bowtie2 and SAMtools. The remaining reads were aligned to the human genome (hg38) using STAR, and PCR duplicates were removed using UMI information with bam-suppress-duplicates. Quality control statistics were generated using fp-framing. Read counting and normalization were performed using fp-count and DESeq2, respectively.

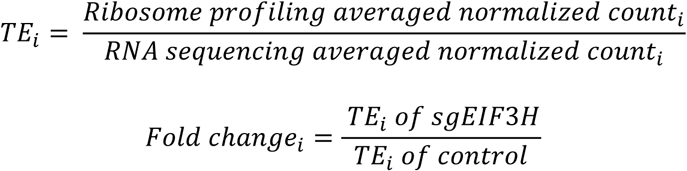

TE and fold-change values were calculated using the averaged normalized counts from biological replicates.

Statistical significance of TE changes was evaluated using the likelihood ratio test implemented in DESeq2.

